# Self-oligomerization regulates stability of Survival Motor Neuron (SMN) protein isoforms by sequestering an SCF^Slmb^ degron

**DOI:** 10.1101/078337

**Authors:** Kelsey M. Gray, Kevin A. Kaifer, David Baillat, Ying Wen, Thomas R. Bonacci, Allison D. Ebert, Amanda C. Raimer, Ashlyn M. Spring, Sara ten Have, Jacqueline J. Glascock, Kushol Gupta, Gregory D. Van Duyne, Michael J. Emanuele, Angus I. Lamond, Eric J. Wagner, Christian L. Lorson, A. Gregory Matera

**Author notes:** Address correspondence to: A. Gregory Matera, Integrative Program for Biological and Genome Sciences, Campus Box 7100, University of North Carolina, Chapel Hill, NC 27599, Voice.

## Abstract

Spinal muscular atrophy (SMA) is caused by homozygous mutations in human *SMN1*. Expression of a duplicate gene (*SMN2*) primarily results in skipping of exon 7 and production of an unstable protein isoform, SMNΔ7. Although *SMN2* exon skipping is the principal contributor to SMA severity, mechanisms governing stability of SMN isoforms are poorly understood. We used a *Drosophila* model system and label-free proteomics to identify the SCF^Slmb^ ubiquitin E3 ligase complex as a novel SMN binding partner. SCF^Slmb^ interacts with a phospho-degron embedded within the human and fruitfly SMN YG-box oligomerization domains. Substitution of a conserved serine (S270A) interferes with SCF^Slmb^ binding and stabilizes SMNΔ7. SMA-causing missense mutations that block multimerization of full-length SMN are also stabilized in the degron mutant background. Overexpression of SMNΔ7^S270A^, but not wild-type SMNΔ7, provides a protective effect in SMA model mice and human motor neuron cell culture systems. Our findings support a model wherein the degron is exposed when SMN is monomeric, and sequestered when SMN forms higher-order multimers.

## Introduction

Spinal muscular atrophy (SMA) is a common neuromuscular disorder, recognized as the most prevalent genetic cause of early childhood mortality (Pearn 1980). Patients with the most severe form of the disease, which is also the most common, become symptomatic in the first six months of life and rarely live past two years (Wee et al. 2010; Prior 2010). Because the onset of symptoms and their severity can vary, SMA has historically been classified into three subtypes (Ogino and Wilson 2004). More recently, clinicians have recognized that SMA is better characterized as a continuous spectrum disorder, ranging from acute (prenatal onset) to nearly asymptomatic (Tiziano et al. 2013). Clinically, SMA patients experience degeneration of motor neurons in the anterior horn of the lower spinal cord (Crawford and Pardo 1996). This leads to progressive atrophy of proximal muscle groups, ultimately resulting in loss of motor function and symmetrical paralysis. The cause of death is often restrictive respiratory failure (Kolb and Kissell 2015).

SMA typically results from homozygous deletion of the *survival motor neuron 1* (*SMN1*) gene (Lefebvre et al. 1995). A small fraction of SMA patients have lost one copy of *SMN1* and the remaining copy contains a point mutation (Burghes and Beattie 2009). Humans have two *SMN* paralogs, named *SMN1* and *SMN2*, both of which contribute to total cellular levels of SMN protein. *SMN2* exon 7 contains a silent base change that alters splicing to primarily produce a truncated, unstable protein product called SMN∆7 (Lorson et al. 1999; Monani et al. 1999; Lorson and Androphy 2000). The last 16 amino acids of SMN are replaced in SMNΔ7 by four amino acids, EMLA, encoded by exon 8. Current estimates suggest that *SMN2* produces 10-15% of the level of full-length protein produced by *SMN1* (Lorson et al. 2010). Complete loss of SMN is lethal in all organisms investigated to date (O’Hearn et al. 2016). Although the amount of full-length protein produced by *SMN2* is not enough to compensate for loss of *SMN1*, *SMN2* is sufficient to rescue embryonic lethality (Monani et al. 2000). SMA is therefore a disease that arises due to a hypomorphic reduction in SMN levels (Lefebvre et al. 1995). Furthermore, relative levels of the SMN protein correlate with the phenotypic severity of SMA (Coovert et al. 1997; Lefebvre et al. 1997).

Whereas a causative link between *SMN1* and SMA was established over 20 years ago, the molecular role of SMN in disease etiology remains unclear. SMN is the central component of a multimeric protein assemblage known as the SMN complex (Matera and Wang 2014; Li et al. 2014). The best-characterized function of this complex, which is found in all tissues of metazoan organisms, is in the cytoplasmic assembly of small nuclear ribonucleoproteins (snRNPs), core components of the spliceosome (Fischer et al. 1997; Meister et al. 2001; Pellizzoni et al. 2002).

Although it is ubiquitously expressed, SMN has also been implicated in a number of tissue-specific processes related to neurons and muscles. These functions include actin dynamics (Oprea et al. 2008; Ackermann et al. 2013), axonal pathfinding (Fan and Simard 2002; McWhorter et al. 2003; Sharma et al. 2005), axonal transport of β-actin mRNP (Rossoll et al. 2003), phosphatase and tensin homolog-mediated (PTEN-mediated) protein synthesis pathways (Ning et al. 2010), translational regulation (Sanchez et al. 2013), neuromuscular junction formation and function (Chan et al. 2003; Kariya et al. 2008; Kong et al. 2009; Voigt et al. 2010), myoblast fusion (Shafey et al. 2005) and maintenance of muscle architecture (Rajendra et al. 2007; Walker et al. 2008; Bowerman et al. 2009).

Ubiquitylation pathways have been shown to regulate the stability and degradation of SMN (Chang et al. 2004; Burnett et al. 2009; Hsu et al. 2010) as well as axonal and synaptic stability (Korhonen and Lindholm 2004). In the ubiquitin proteasome system (UPS), proteins destined for degradation are tagged by linkage to ubiquitin through the action of three factors (Petroski 2008). E1 proteins activate ubiquitin and transfer it to the E2 enzyme. E2 proteins conjugate ubiquitin to their substrates. E3 proteins recognize the substrate and assist in the transfer of ubiquitin from the E2. Because E3 ligases confer substrate specificity, they are typically considered as candidates for targeted inhibition of protein degradation. Ubiquitin homeostasis is thought to be particularly important for neuromuscular pathology in SMA (Groen and Gillingwater 2015). Indeed, mouse models of SMA display widespread perturbations in UBA1 (ubiquitin-like modifier activating enzyme 1) levels (Wishart et al. 2014). Furthermore, mutations in UBA1 are known to cause X-linked infantile SMA (Ramser et al. 2008; Schmutzler et al. 2008).

Given the importance of these processes to normal development as well as neurodegenerative disease, we set out to identify and characterize novel SMN binding partners. Previously, we developed *Drosophila melanogaster* as a model system wherein the endogenous *Smn* gene is replaced with a *Flag-Smn* transgene (Praveen et al. 2012). Although it is highly similar to human *SMN1* and *SMN2*, the entire open reading frame of fruitfly *Smn* is contained within a single exon, and so only full-length SMN protein is expressed in *Drosophila* (Rajendra et al. 2007). When modeled in the fly, SMA-causing point mutations recapitulate the full range of phenotypic severity seen in humans (Praveen et al. 2014; Garcia et al. 2016). Using this system, we carried out proteomic profiling of Flag-purified embryonic lysates and identified the SCF^Slmb^ E3 ubiquitin ligase complex as a novel SMN interactor. Importantly, this interaction is conserved from flies to humans. We show that SCF^Slmb^ binding requires a phospho-degron motif located within the SMN self-oligomerization domain, mutation of which stabilizes SMN∆7 and, to a lesser extent, full-length SMN. Additional studies in flies, mice and human cells elucidate a disease-relevant mechanism whereby SMN protein stability is regulated by self-oligomerization. Other E3 ligases have been reported to target SMN for degradation in cultured human cells (Han et al. 2016; Hsu et al. 2010; Kwon et al. 2013). Given our findings in fruit fly embryos, SMN is likely targeted by multiple E3 ubiquitin ligases.

## Results

### Flag-SMN interacts with UPS (ubiquitin proteasome system) proteins

We previously generated transgenic flies that express Flag-tagged SMN proteins in an otherwise null *Smn* background (Praveen et al. 2012). To preserve endogenous expression patterns, the constructs are driven by the native promoter and flanking sequences. As described in the Methods, we intercrossed hemizygous *Flag-Smn*^*WT*^,*Smn*^*X7*^/*Smn*^*D*^ animals to establish a stock wherein all of the SMN protein, including the maternal contribution, is epitope-tagged. After breeding them for >100 generations, essentially all of the animals are homozygous for the *Flag-Smn*^*WT*^ transgene, but second site recessive mutations are minimized due to the use of two different *Smn* null alleles. Adults from this stock display no apparent defects and have an eclosion frequency (~90%) similar to that of wild-type (Oregon-R) animals.

We collected (0-12h) embryos from *Flag-Smn*^*WT/WT*^,*Smn*^*X7/D*^ (SMN) and Oregon-R (Ctrl) animals and analyzed Flag-purified lysates by ‘label-free’ mass spectrometry. In addition to Flag-SMN, we identified SMN complex components Gemin2 and Gemin3, along with all seven of the canonical Sm-core snRNP proteins (Fig. 1A). We also identified the U7-specific Sm-like heterodimer Lsm10/11 (Pillai et al. 2003) and the Gemin5 orthologue, Rigor mortis (Gates et al. 2004). Previous studies of Schneider2 (S2) cells transfected with epitope-tagged *Smn* had identified most of the proteins listed above as SMN binding partners in *Drosophila* (Kroiss et al. 2008). However, despite bioinformatic and cell biological data indicating that Rigor mortis is part of the fruit fly SMN complex, this protein failed to co-purify with SMN in S2 cells (Kroiss et al. 2008; Cauchi et al. 2010; Guruharsha et al. 2011). On the basis of our purification data, we conclude that the conditions are effective and that Rigor mortis/Gemin5 is an integral member of the SMN complex in flies.

**Figure 1:**
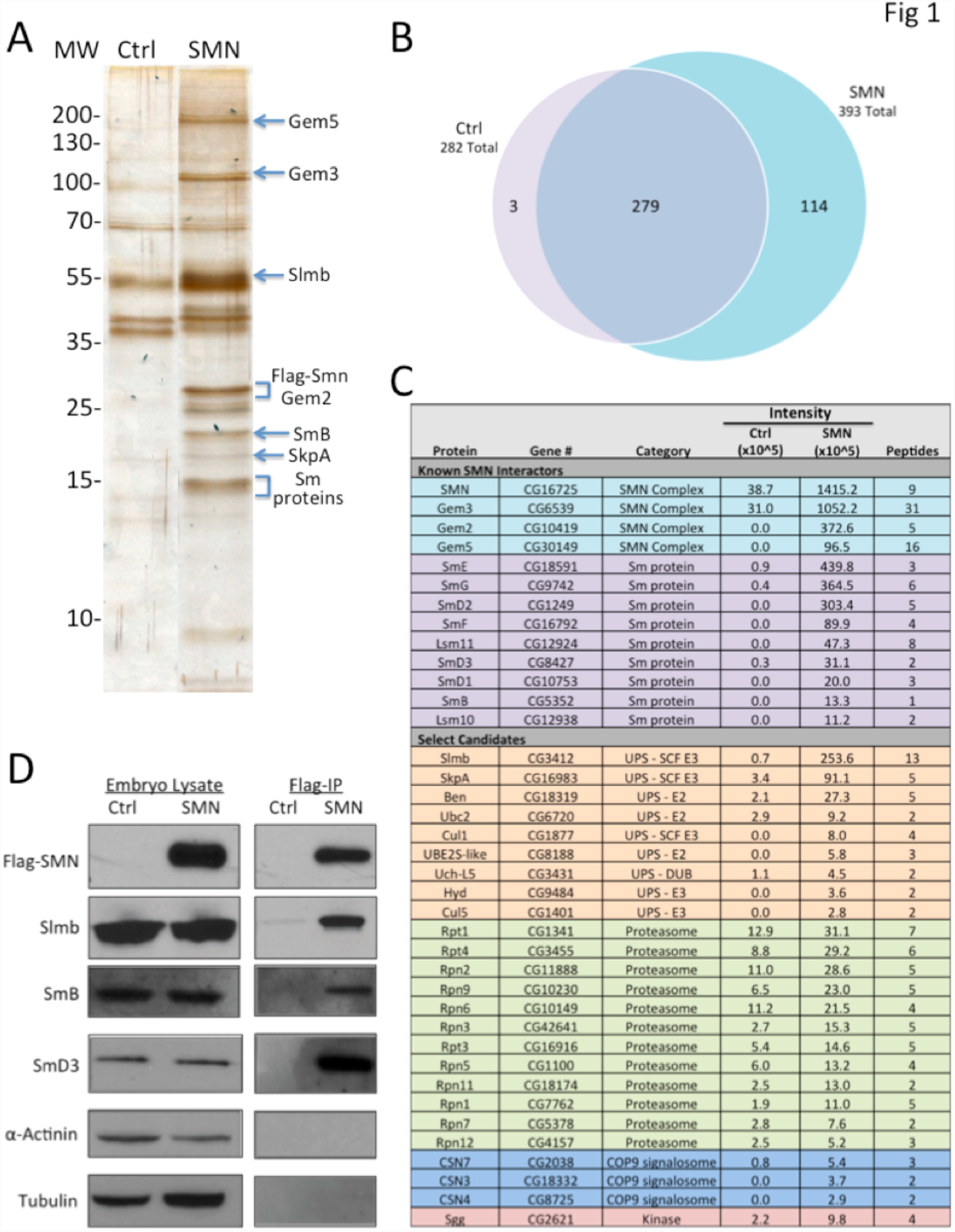
Flag-SMN immunopurified lysates contain known protein interaction partners and ubiquitin proteasome system (UPS) proteins ***A**.* Lysates from Oregon-R control (Ctrl) *Drosophila* embryos and embryos expressing only transgenic Flag-SMN (SMN) were Flag-immunopurified and protein eluates were separated by gel electrophoresis and silver stained. Band identities predicted by size using information from panels C and D. ***B***. Direct mass spectrometric analysis of the eluates (which were not gel purified) identified a total of 396 proteins, 114 of which were detected only in SMN sample and 279 of which were detected in both SMN and Ctrl samples. ***C***. Flag-purified eluates were analyzed by ‘label-free’ mass spectrometry. Numerous proteins that copurify with Flag-SMN are part of the ubiquitin proteasome system (UPS). Of these UPS proteins, Cullin 1 (Cul 1), SkpA, and supernumerary limbs (Slmb), were highly enriched (at least 10 fold) in the SMN sample as compared to Ctrl. ***D***. Western blot analysis of Flag-purified eluates was used to further validate the presence or absence of SMN interaction partners. Flag-SMN was successfully purified from SMN embryos, but was undetectable in the control. As positive controls for known protein interaction partners of SMN, SmB and SmD3 were also easily detectable by western blotting using anti-Sm antibodies. The presence of Slmb was verified using anti-SImb. α-Actinin and Tubulin were not enriched in our purification and are used as negative controls to demonstrate specificity.

A detailed proteomic analysis of these flies will be presented elsewhere. As shown in Fig. 1B, our preliminary analysis identified 396 proteins, 114 of which were detected only in the Flag-SMN sample and not in the control. An additional 279 proteins were detected in both the Flag purification and control samples. In addition to SMN complex members, we co-purified numerous factors that are part of the ubiquitin proteasome system (UPS; Fig. 1C). Among these UPS proteins, we identified Cullin 1 (Cul1), Skp1-related A (SkpA), and supernumerary limbs (Slmb), as being highly enriched (>10 fold) in Flag-SMN samples as compared to the control. Together, these proteins comprise the SCF^Slmb^ E3 ubiquitin ligase. Cul1 forms the major structural scaffold of this horseshoe-shaped, multi-subunit complex (Zheng et al. 2002). Slmb is a F-box protein and is the substrate recognition component (Jiang and Struhl 1998). SkpA is a bridging protein essential for interaction of Cul1 with the F-box protein (Patton et al. 1998a; Patton et al. 1998b). Because of its role in substrate recognition, Slmb is likely to be the direct interacting partner of SMN within the SCF^Slmb^ complex. For this reason, we focused on Slmb for the initial validation. As shown, Slmb was easily detectable in Flag-purified eluates from embryos expressing Flag-SMN and nearly undetectable in those from control embryos (Fig. 1D). SmB and SmD3 were also easily detectable by western blot in Flag-purified embryonic lysates and were used as positive controls for known protein interaction partners of SMN. Tubulin and α-Actinin were not detected as interacting with SMN in our purification and demonstrate the specificity of the detected SMN interactions.

### SCF^Slmb^ is a *bona fide* SMN interaction partner that ubiquitylates SMN

As an E3 ubiquitin ligase, the SCF^Slmb^ complex is a substrate recognition component of the ubiquitin proteasome system. As outlined in Fig. 2A, E3 ligases work with E1 and E2 proteins to ubiquitylate their targets. The interaction of SCF^Slmb^ with SMN was verified in a reciprocal co-immunoprecipitation, demonstrating that Flag-tagged SCF components form complexes with endogenous SMN (Fig. 2B) in S2 cells.

**Figure 2:**
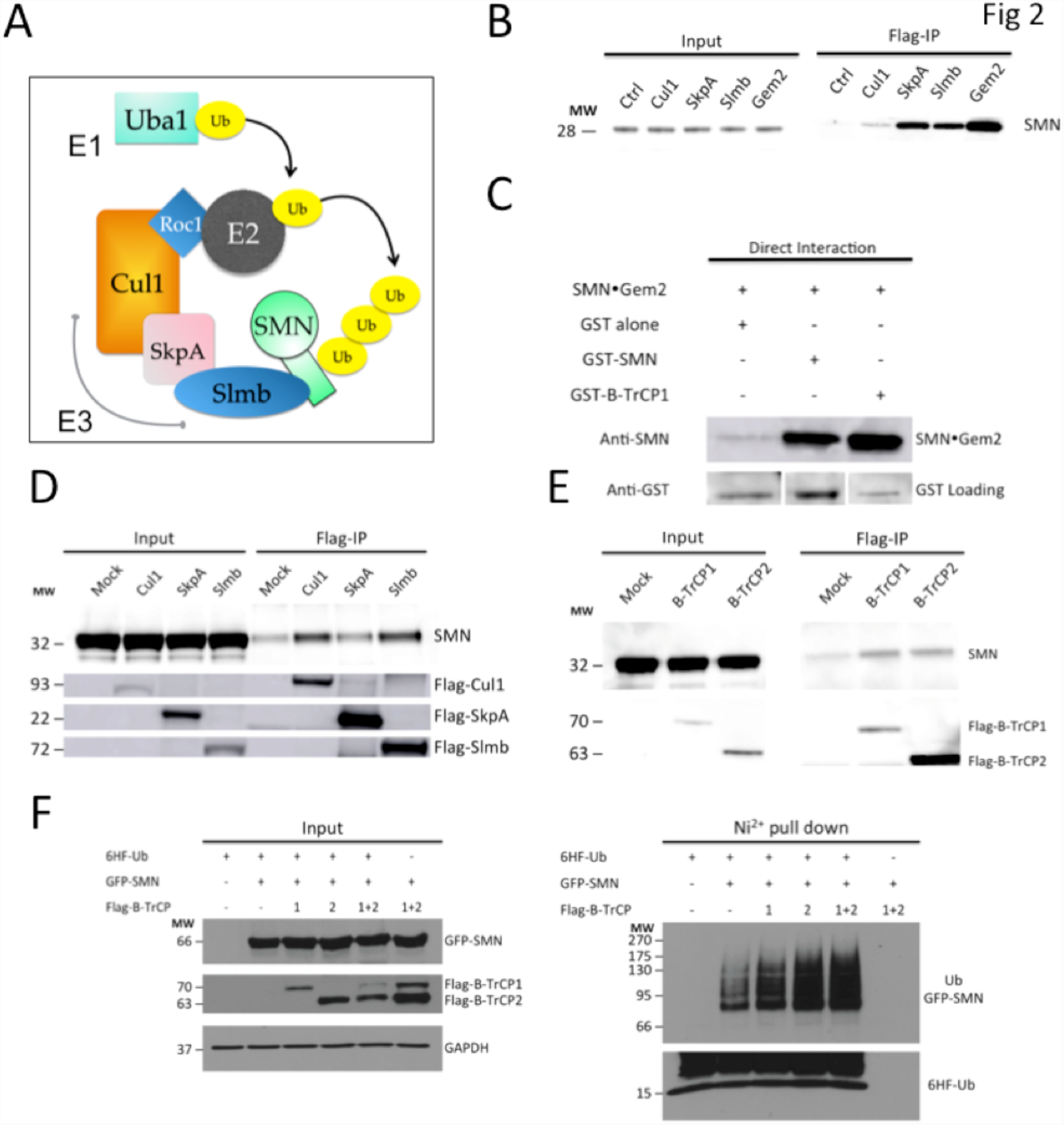
Conserved interaction between SMN and the SCF^slmbB-TrCP^ E3 ubiquitin ligase results in ubiquitylation of SMN. ***A***. E3 ligases work with E1 and E2 proteins to ubiquitylate their targets. The SCF^slmbB-TrCP^ E3 ubiquitin ligase is made up of three proteins: Cul1, SkpA, and Slmb. The E3 ubiquitin ligase is the substrate recognition component of the ubiquitin proteasome system. ***B***. Following Cul1-Flag, SkpA-Flag, Flag-Slmb and Flag-Gem2 immunoprecipitation from Drosophila S2 cell lysates, western analysis using anti-SMN antibody for endogenous SMN was carried out. Co-purification of each of the SCF components with endogenous SMN was detected. ***C.*** An *in vitro* binding assay tested direct interaction between human SMN∆5-Gemin2 (SMN•Gem2) (Martin et al. 2012; Gupta et al. 2015) and purified GST-tagged proteins. SMN•Gem2 did not interact with GST protein alone, but bound to GST tagged *Drosophila* SMN (GST-SMN) and GST tagged human B-TrCP1 (GST-B-TrCP1). Levels of GST alone, GST-SMN, and GST-B-TrCP1 were detected using anti-GST antibody. ***D***. The interaction of Flag-tagged Drosophila SCF components with endogenous human SMN was tested in in HEK 293T cells. Human SMN was detected at high levels following immunoprecipitation of Drosophila Flag-Cul1 and Flag-Slmb and detected at a lower level following Drosophila Flag-SkpA immunoprecipitation. ***E.*** Flag-tagged versions of the human homologs of Slmb, Flag-B-TrCP1 and Flag-B-TrCP2, interact with endogenous human SMN in HEK 293T cells demonstrated by Flag-immunopurification followed by immunodetection of SMN. ***F***. Protein lysate from HEK 293T cells transfected with 6xHis-Flag-ubiquitin (6 HF-Ub) and GFP-SMN was purified using a Ni^2+^ pu11 down for the tagged ubiquitin. Baseline levels of ubiquitylated GFP-SMN were detected using anti-GFP antibody. Following transfection of Flag-B-TrCP1 or Flag-B-TrCP2, the levels of ubiquitylated SMN markedly increased. Ubiquitylation levels were further increased following addition of both proteins together. In the input, GFP-SMN was detected using anti-GFP antibody, Flag-B-TrCP1 and Flag-B-TrCP2 were detected using anti-Flag antibody, and GAPDH was detected by anti-GAPDH antibody. In the Ni^2+^ pull down, ubiquitylated GFP-SMN was detected using anti-GFP antibody and 6HF-Ub was detected using anti-Flag antibody to verify successful pull down of tagged ubiquitin.

SCF complexes are highly conserved from flies to humans: SkpA is 77% identical to human Skp1, Cul1 is 63% identical, and Slmb is 80% identical to its human homologs, B-TrCP1 and B-TrCP2. Slmb/B-TrCP is the SCF component that directly contacts substrates of the E3 ligase. We therefore tested the interaction of purified human SMN in complex with Gemin2 (SMN•Gem2) (Martin et al. 2012; Gupta et al. 2015) with GST-B-TrCP1 in an *in vitro* binding assay (Fig. 2C). SMN•Gem2 did not interact with GST alone, but was detected at high levels following pulldown with either the positive control *Drosophila* GST-SMN or GST-B-TrCP1. We also tested the interaction of Flag-tagged *Drosophila* SCF components with endogenous human SMN in HEK 293T cells (Fig. 2D). Accordingly, human SMN was co-precipitated with Flag-Cul1 and Flag-Slmb and at lower levels following Flag-SkpA immunoprecipitation. Flag-B-TrCP1 and Flag-B-TrCP2, the two human homologs of Slmb, also copurified with endogenous human SMN in HEK 293T cells (Fig. 2E). Altogether, these data demonstrate a conserved interaction between SMN and the SCF^Slmb/B-TrCP^ E3 ubiquitin ligase complex.

In order to test the functional consequences of this conserved interaction between SMN and SCF^Slmb/B-TrCP^, a cell based ubiquitylation assay was performed (Fig. 2F). Protein lysate from HEK 293T cells transfected with 6xHis-Flag-ubiquitin and GFP-SMN was purified using a Ni^2+^ pull down for the tagged ubiquitin. Baseline levels of ubiquitylated GFP-SMN were detected using anti-GFP antibody. Following transfection of Flag-B-TrCP1 or Flag-B-TrCP2, the levels of ubiquitylated SMN markedly increased (Fig. 2F). Ubiquitylation levels were further increased following addition of both proteins together. These experiments demonstrate that SCF^Slmb/B-TrCP^ can ubiquitylate SMN *in vivo*.

### Depletion of Slmb/B-TrCP results in a modest increase in SMN levels

Given that one of the primary functions of protein ubiquitylation is to target proteins to the proteasome, we examined whether depletion of Slmb by RNA interference (RNAi) using dsRNA in S2 cells would increase SMN levels (Fig. 3A). Following Slmb RNAi, endogenous SMN levels were modestly increased as compared to cells treated with control dsRNA. We obtained similar results using an siRNA that targets both B-TrCP1 and B-TrCP2 in HeLa cells. As shown in Fig. 3B, we detected a modest increase in levels of full-length SMN following B-TrCP RNAi, but not control RNAi. Next, we treated S2 cells with cycloheximide (CHX), in the presence or absence of dsRNA targeting Slmb, to determine whether differences in SMN levels would be exacerbated when production of new proteins was prevented (Fig. 3C). SMN protein levels were also directly targeted using dsRNA against *Smn* as a positive control for the RNAi treatment. At 6 hours post-CHX treatment there was a modest increase in full-length SMN levels following Slmb RNAi as compared to the initial timepoint (0h) or the negative control (Ctrl) RNAi (Fig. 3C). Together, these data indicate that Slmb/B-TrCP participates in the regulation of SMN protein levels.

**Figure 3:**
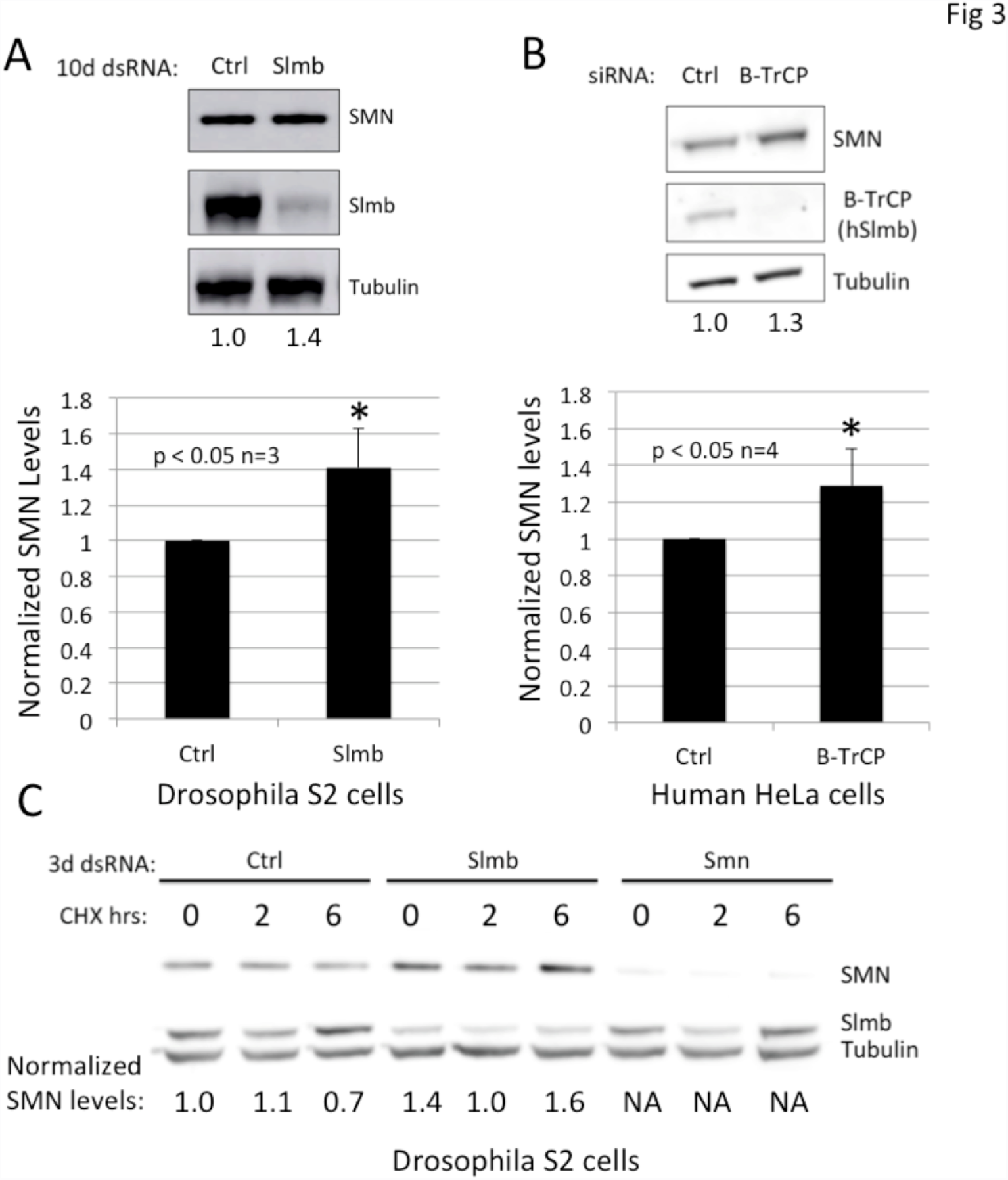
Depletion of Slmb/B-TrCP results in an increase of SMN levels. ***A.*** Depletion of Slmb using 10 day (1 Od) treatment with dsRNA in *Drosophila* S2 cells resulted in modestly increased SMN levels. Following Slmb RNAi, full-length SMN levels were increased as compared to cells treated with control dsRNA against Gaussia Luciferase, which is not expressed in S2 cells. ***B.*** The effect of B-TrCP depletion on SMN levels in human cells was tested using siRNA that targets both B-TrCP1 and B-TrCP2 in HeLa cells. We detected a modest increase in levels of full-length endogenous SMN after B-TrCP RNAi but not control (scramble) RNAi. ***C.*** *Drosophila* S2 cells were treated with cycloheximide (CHX), an inhibitor of protein synthesis, following Slmb depletion following a 3d dsRNA treatment to test whether differences in protein levels would be exacerbated when the production of new protein was prevented. SMN protein levels were also directly targeted using dsRNA against *Smn* as a positive control for the RNAi treatment. As a negative control (Ctrl), dsRNA against *oskar*, which is not expressed in S2 cells, was used. Protein was collected at 0, 2, and 6 hours post CHX treatment. At 6 hours post-CHX treatment there is a modest increase in full-length SMN levels following *Slmb* RNAi as compared to the initial timepoint (0h) and as compared to control RNAi treatment.

### Identification and characterization of a Slmb/B-TrCP degradation signal in SMN

Studies of numerous UPS substrates in a variety of species have revealed the presence of degradation signals (called degrons) that are required for proper E3 target recognition and binding. Slmb/B-TrCP canonically recognizes a consensus DpSGXXpS/T degron, where p indicates a phosphoryl group (Jin et al. 2005; Frescas and Pagano 2008; Fuchs et al. 2004). There are also several known variants of this motif, for example: DDGFVD, SSGYFS, TSGCSS (Kim et al. 2015). As shown in Fig. 4A, we identified a putative Slmb/B-TrCP degron (^269^MSGYHT^274^) in the highly conserved self-oligomerization domain (YG Box) of human SMN. Interestingly, this sequence was previously identified as part of a larger degron motif (^268^YMSGYHTGYYMEMLA^282^) that was thought to be created in SMN∆7 by *SMN2* alternative splicing (Cho and Dreyfuss 2010). In particular, mutation of S270 (S201 in flies) to alanine was shown to dramatically stabilize SMN∆7 constructs in human cells, and overexpression of SMN∆7^S270A^ in SMN-deficient chicken DT40 cells rescued their viability (Cho and Dreyfuss 2010). However, factors responsible for specifically mediating SMN∆7 degradation have not been identified.

**Figure 4:**
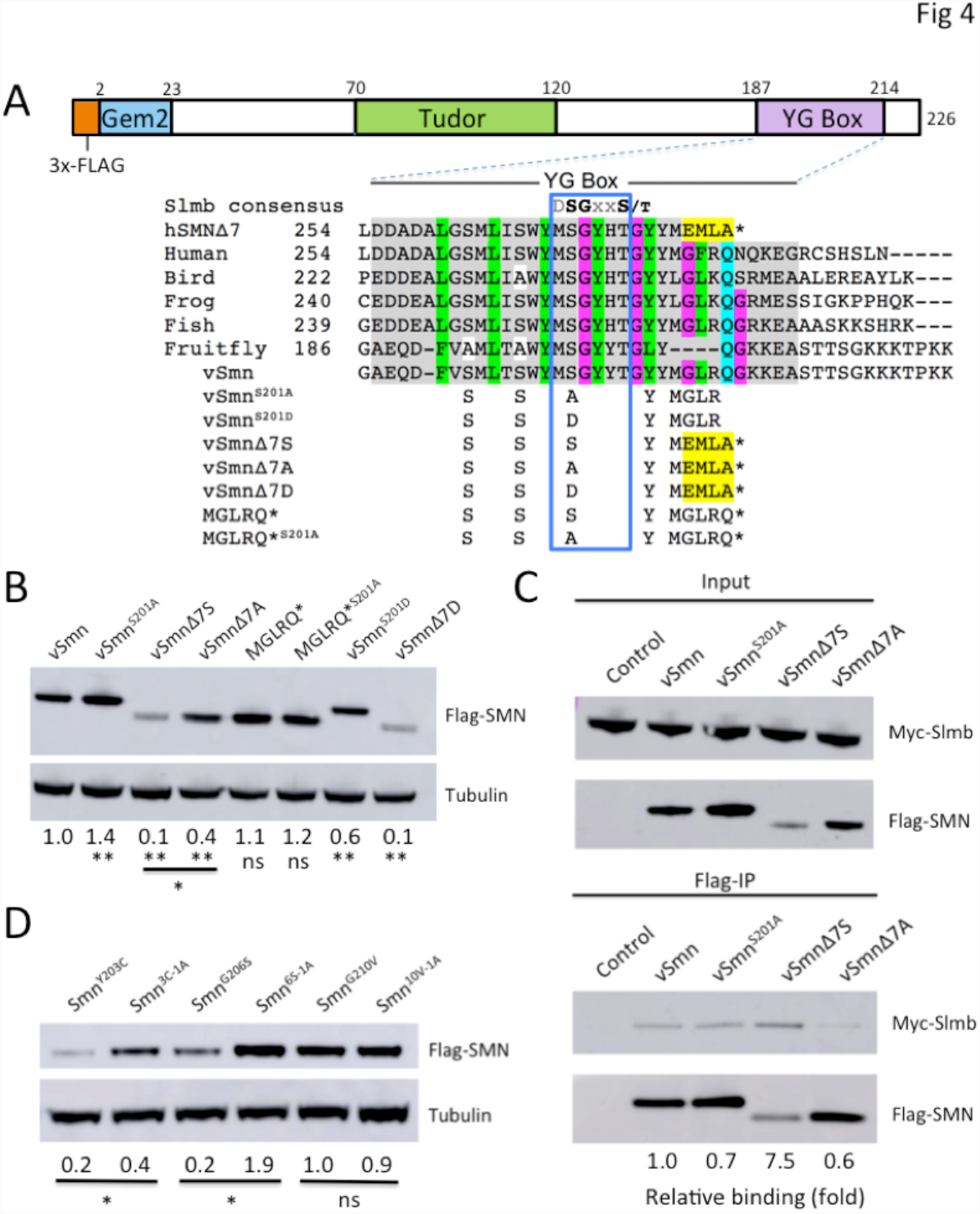
Identification and mutation of a putative Slmb/B-TrCP phospho-degron ***A.*** Identification of a conserved putative Slmb phospho-degron (DpSGXXpS/T motif variant) in the C-terminal self-oligomerization domain (YG Box) of SMN. The amino acid sequence of Smn from a variety of vertebrates is shown to illustrate conservation of this motif and rationale for the amino acid changes. Full-length human SMN is labeled as “Human” and the truncated isoform is labeled “hSMN∆7”. Endogenous *Drosophila melanogaster* Smn is labeled “Fruitfly”. To generate a more vertebrate-like SMN, key amino acids in Drosophila SMN were changed to amino acids conserved in vertebrates. Using this SMN backbone, a serine to alanine mutation was made in the putative degron in both full-length (vSMN^S201A^) and truncated SMN∆7 (vSMN∆7A). An additional SMN construct that is the same length as SMN∆7, but has the amino acid sequence GLRQ (the next amino acids in the sequence) rather than EMLA (the amino acids introduced by mis-splicing of *SMN2*) was made. The same serine to alanine mutation was made in this construct as well (MGLRQ* and MGLRQ*^S201A^). Finally, to mimic a phosphorylated serine the full-length SMN construct (vSmn^S201D^) and truncated SM N (vSMN∆7D) were made. ***B.*** Western blotting was used to determine protein levels of each of these SMN constructs, with expression driven by the endogenous promoter, in Drosophila S2 cells. Both the vSMN and vSMN∆7S proteins show increased levels when the serine is mutated to an alanine, indicating disruption of the normal degradation of SMN. Additionally, MGLRQ* protein is present at higher levels than is vSMN∆7S and protein levels do not change when the serine is mutated to an alanine. Normalized fold change as compared to vSmn levels is indicated at the bottom. *p<0.05, **p<0.01 n=3. ***C.*** Flag-tagged SMN constructs were co-transfected with Myc-Slmb in Drosophila S2 cells. Protein lysates were Flag-immunoprecipitated and probed with anti-Myc antibody to detect SMN-Slmb interaction. In both full-length SMN (vSMN) and truncated SMN (vSMN∆7), serine to alanine mutation decreased interaction of Slmb with SMN. Truncated SMN (vSMN∆7) showed a dramatically increased interaction with Slmb as compared to full-length SMN (vSMN), despite the fact it is present at lower levels. ***D.*** Full-length SMN constructs containing point mutations known to decrease self-oligomerization (Smn^Y203C^ and Smn^G206S^) and a mutation that does not disrupt self-oligomerization in the fly (Smn^G210V^) with and without the serine to alanine mutation were transfected in Drosophila *S2* cells. The constructs containing the serine to alanine mutation are as follows: Smn^Y203C^->Smn^3C-1A^, Smn^G2065^->Smn^6S-1A^, Smn^G210V^->Smn^l0V-1A^. The serine to alanine mutation has a stabilizing effect on SMN mutants with poor self-oligomerization capability. *p<0.05, n=3.

**Figure 5:**
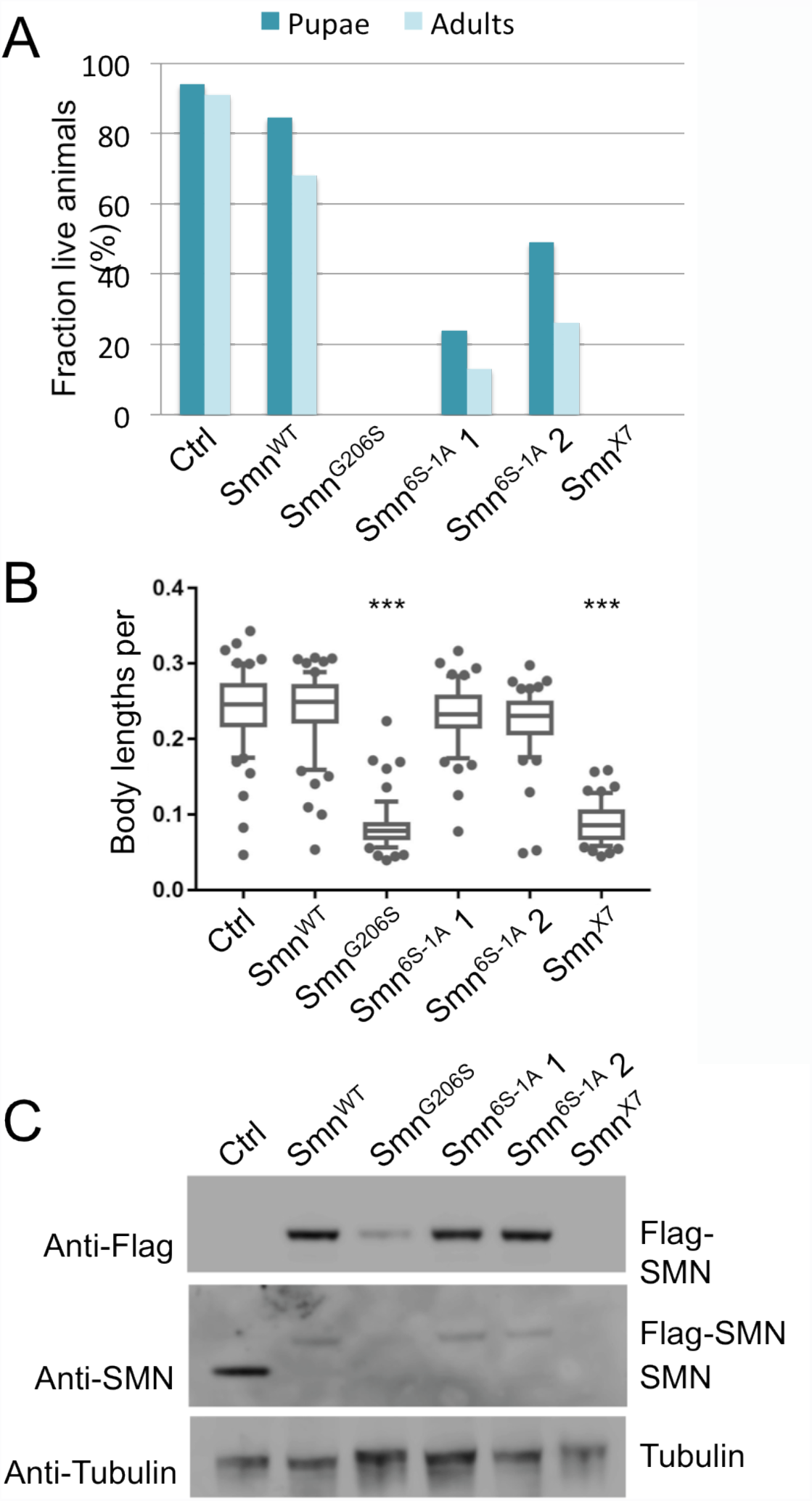
Mutation of the SImb degron rescues defects in SMA model flies. ***A.*** Viability analysis of an SMA point mutation (G206S) in the presence and absence of the degron mutation, S201A. Flies with the following genotypes were analyzed in this experiment: Oregon-R (Ctrl), *Flag-Smn*^*V/T*^,*Smn*^*X7*^/*Smn*^*X7*^ (Smn^WT^), *Flag-Smn*^*G206S*^,*Smn*^*X7*^/*Snn*^*X7*^ (Smn^G2ces^), *Flag-Smn*^*G206,S201A*^,*Smn*^*X7*^/*Smn*^*X7*^ (Smn^6S-1A^) or *Smn*^*X7*^/*Smn*^*X7*^ (Smn^X7^). The data for each genotype are expressed as a fraction of pupae or adults over the total number of starting larvae, n=200. Expression of the WT transgene (*Smn*^WT^) shows robust rescue of the null (*Smn*^*X7*^) phenotype (∼68% adults). *Smn*^G206S^ is a larval lethal mutation. In two independent recombinant lines of *Smn*^*6S-1A*^ (*Smn*^*6S-1A*^1 and *Smn*^*6S-1A*^2) a fraction of the larvae complete development to become adults. *B.* Locomotor ability of early 3^rd^ instar larvae was determined by tracking their movement for one minute and then calculating the velocity. To account for potential differences in larval size, speed is expressed as average body lengths per second moved. Genotypes are as in panel A. *Smn*^*G206S*^ larvae move similarly to null animals. The motility of *Smn*^*6S-1A*^*1* and *Smn*^*6S-1A*^*2* larvae is not different from Ctrl or *Smn*^*WT*^ larvae. ***p<0.001, n=50 to 60 larvae. *C.* Larval protein levels were examined by western blotting; genotypes as in panel A. Lysates from hemizygous mutant lines were probed with anti-Flag or anti-SMN antibodies as indicated. The slower migrating bands represent the Flag-tagged transgenic proteins and the faster migrating band corresponds to endogenous SMN, which is present only in the Ctrl (note Oregon-R has two copies *Smn* whereas the transgenicshave only one). *Smn*^*G206S*^ has very low levels of SMN protein. Flies bearing the S201A degron mutation in addition to G206S (*Smn*^*6S-1A*^) expressmarkedly increased levels of SMN protein.

In order to develop a more disease-relevant *Drosophila* system to investigate SMN YG box function, we generated a ‘vertebrate-like’ SMN construct, called vSmn (Fig. 4A). Transgenic flies expressing Flag-vSmn and Flag-vSmn^S201A^ in the background of an *Smn*^*X7*^ null mutation are fully viable (Fig. S1). In fact, the eclosion frequencies of these animals are consistently higher than those that express Flag-Smn^WT^ (Fig. S1). Additional *Smn* mutant constructs were generated using the vSmn backbone, including both the full-length (e.g. vSmn^S201A^) and truncated (e.g. vSmnΔ7A) versions of the protein (Fig. 4A). To test the effects of overall protein length and distance of the putative degron from the C-terminus, we also generated vSmn constructs that are the same length as SMNΔ7, replacing the MEMLA* (the amino acids introduced by human *SMN2* splicing) with MGLRQ*, see Fig. 4A. The S201A mutation was created in this construct as well (MGLRQ*^S201A^). To mimic a constitutively phosphorylated state, we also introduced serine to aspartate mutations, vSmn^S201D^ and vSmn∆7D. We transfected each of these constructs, Flag-tagged and driven by the native *Smn* promoter, into S2 cells and measured protein levels by western blotting (Fig. 4B). We note that these constructs are expressed at levels far below those of endogenous SMN protein in S2 cells; moreover, they do not affect levels of endogenous SMN (Fig. S2). As shown, the vSmn^S201A^ and vSMNΔ7A constructs exhibited increased protein levels compared to their serine containing counterparts, whereas levels of the S201D mutants were reduced, suggesting that the phospho-degron motif identified in human SMN∆7 (Cho and Dreyfuss 2010) is also conserved in the fly protein. In addition to examining protein levels of each of these constructs in cell culture, transgenic flies expressing vSmn, vSmn^S201A^, vSmnΔ7S, and vSmnΔ7A were created. Here again we observed that S201A mutation increased protein levels of both full-length SMN and SMN∆7 (Fig. S3).

The MGLRQ* construct is present at levels that are similar to wild-type (vSmn) and much higher than vSmnΔ7S. Based on the crystal structures of the SMN YG box (Martin et al. 2012; Gupta et al. 2015), the presence of the MGLR insertion in *Drosophila* SMN is predicted to promote self-oligomerization (A.G. Matera and G.D. Van Duyne, unpublished), thus stabilizing the protein within the SMN complex (Burnett et al. 2009). By the same logic, the relative inability of vSmnΔ7S to self-interact would be predicted to lead to its destruction. To determine whether the observed increase in SMN protein levels correlated with its ability to interact with Slmb, we co-transfected the appropriate Flag-Smn constructs with Myc-Slmb in S2 cells. Protein lysates were then Flag-immunoprecipitated and probed with anti-Myc antibody (Fig. 4C). The S201A mutation decreased binding of Slmb to both the full-length and the truncated SMN isoforms (Fig. 4C). However, the vSmn∆7S construct co-precipitated the greatest amount of Slmb protein, despite the fact that it is present at much lower levels in the input lysate (Fig. 4C). Because SMN∆7 is defective in self-interaction, this result suggests that the degron is more accessible to Slmb when SMN is monomeric and cannot efficiently oligomerize.

### SMN self-oligomerization regulates access to the Slmb degron

To examine the connection between SMN self-oligomerization and degron accessibility more closely we took advantage of two SMA patient-derived point mutations (Y203C and G206S) that are known to destabilize the full-length protein and to decrease its self-oligomerization capacity (Praveen et al. 2014). As a control, we also employed an SMA-causing mutation (G210V) that does not disrupt SMN self-oligomerization (Praveen et al. 2014; Gupta et al. 2015). Next, we introduced the S201A degron mutation into all three of these full-length SMN constructs, transfected them into S2 cells and carried out western blotting (Figs. 4D and S2). The S201A degron mutation has a clear stabilizing effect on the G206S and Y203C constructs, as compared to the effect of S201A paired with G210V. Hence, we conclude that the Slmb degron is exposed when SMN is present predominantly as a monomer, whereas it is less accessible when the protein is able to form higher order multimers.

### Mutation of the Slmb degron rescues viability and locomotion defects in SMA model flies

Next, we examined the effect of mutating the Slmb degron in the context of the full-length protein *in vivo*. We characterized adult viability, larval locomotion and SMN protein expression phenotypes of the G206S mutants in the presence or absence of the degron mutation, S201A (Fig. 5A-C). As described previously (Praveen et al. 2014), *Smn*^*G206S*^ animals express very low levels of SMN and fail to develop beyond larval stages. In contrast, flies bearing the S201A degron mutation in addition to G206S (*Smn*^*6S-1A*^) express markedly increased levels of SMN protein (Fig. 5C) and a good fraction of these animals complete development (Fig. 5A). Moreover, *Smn*^*6S-1A*^ larvae display significantly improved locomotor activity as compared to *Smn*^*G206S*^ or *Smn*^*X7*^ null mutants (Fig. 5B). These results strongly suggest that both the structure of the G206S mutant protein as well as its instability contribute to the organismal phenotype.

### GFP-SMN∆7 overexpression stabilizes endogenous SMN and SMN∆7 in cultured human cells

Increased *SMN2* copy number correlates with a milder clinical phenotype in SMA patients (Oskoui et al. 2016). This phenomenon was successfully modeled in mice over a decade ago (Monani et al. 2000; Hsieh-Li et al. 2000), showing that high copy number *SMN2* transgenes fully rescue the null phenotype, whereas low copy transgenes do not. Moreover, transgenic expression of a human *SMN∆7* cDNA construct in a low-copy *SMN2* background improves survival of this severe SMA mouse model from P5 (post-natal day 5) to P13 (Le et al. 2005). Although the truncated SMN likely retains partial functionality, the protective effect of *SMN∆7* overexpression may not entirely be intrinsic to the protein. That is, SMN∆7 could also act as a ‘soak-off’ factor, titrating the ubiquitylation machinery and stabilizing endogenous SMN. In such a scenario, the prediction would be that SMN∆7A is less protective than SMN∆7S because it is not a very good substrate for SCF^Slmb^.

We therefore compared the stabilizing effects of overexpressing GFP-tagged SMNΔ7^S270A^ (SMN∆7A) and SMNΔ7 (SMN∆7S) on endogenous human SMN and SMN∆7. HEK 293T cells were transfected with equivalent amounts of GFP-SMN∆7A or -SMN∆7S. The following day, cells were harvested after treatment with cycloheximide (CHX) for zero to ten hours. As shown in Fig. 6A, western blotting with anti-SMN showed that the SMN∆7S construct exhibits a clear advantage over SMN∆7A in its ability to stabilize endogenous SMN and SMN∆7. By comparing band intensities within a given lane, we generated average intensity ratios for each time point using replicate blots (Fig. 6A, table). We then calculated a ‘stabilization factor’ by taking a ratio of these two ratios. As shown (Fig. 6A, graph), the protective benefit of overexpressing ∆7S vs. ∆7A at t=0 hr was roughly 3.0x for endogenous SMN∆7 and 1.75x for full-length SMN. Thus, as predicted above, the GFP-SMN∆7A construct was much less effective at stabilizing endogenous SMN isoforms. Because SMN∆7 is a relatively good SCF^Slmb^ substrate, overexpression of this isoform protects full-length SMN from degradation.

**Figure 6:**
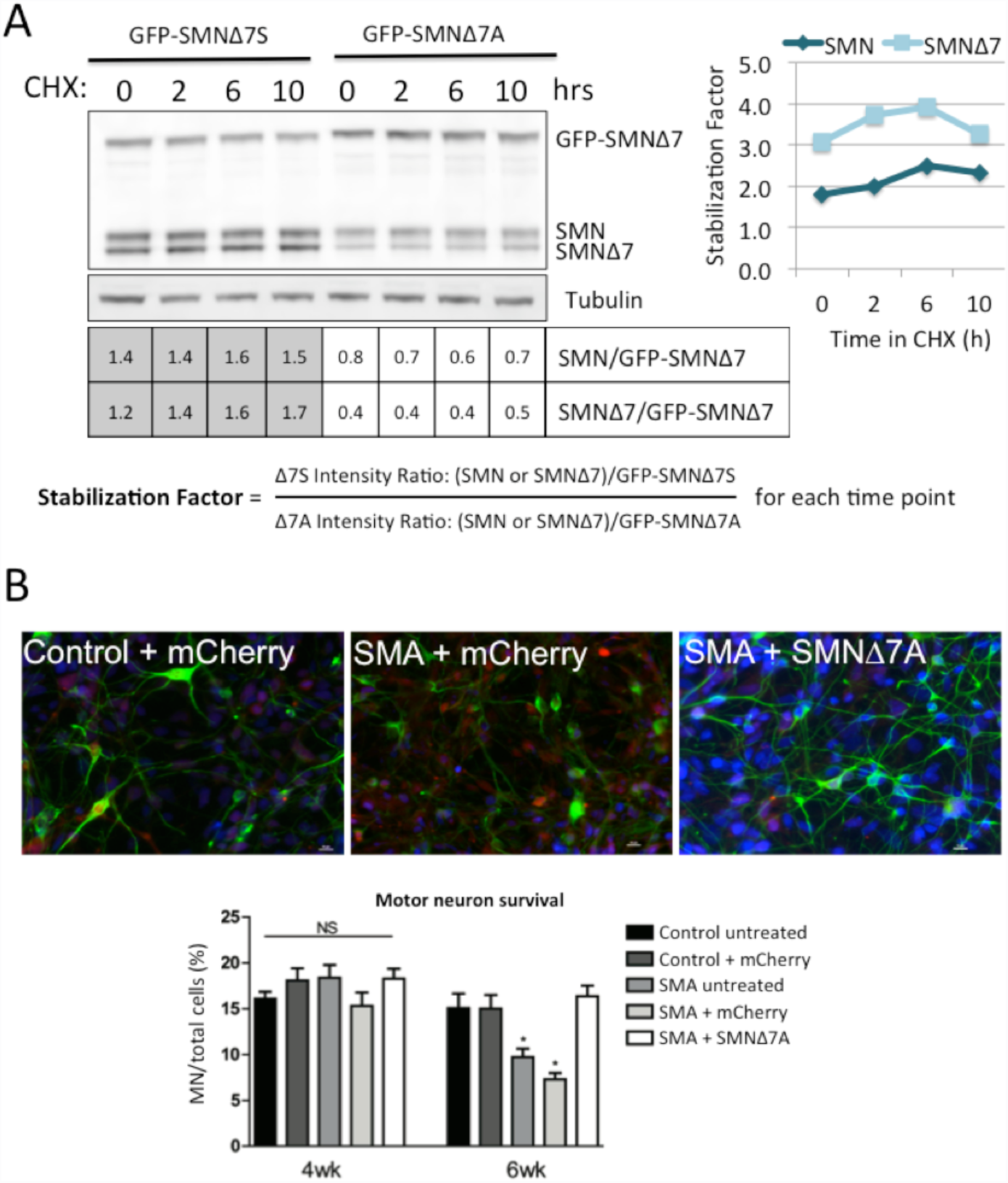
Stabilization of endogenous SMN and SMN∆7 in cultured human cells. *A.* HEK 293T cells were transfected with equivalent amounts of GFP-SMN∆7A or-SMN∆7S. The following day, cells were harvested after treatment with cycloheximide (CHX) for zero to ten hours. Western blotting with anti-SMN showed that SMN∆7S stabilizes endogenous SMN and SMN∆7 to a greater extent than SMN∆7A. By comparing band intensities within a given lane, we generated average intensity ratios for each time point using replicate blots. We then calculated a ‘stabilization factor’ by taking a ratio of these two ratios. The protective benefit of overexpressing A7S vs. Δ7Α at t=0 hr was roughly 3.Ox for endogenous SMN∆7 and 1,75x for full-length SMN. *B.* SMN∆7A (S270A) expression protects SMA iPSC-derived motor neurons. Control motor neurons were left untreated or transduced with a lentiviral vector expressing an mCherry control. SMA motor neurons were left untreated, transduced with a lentiviral vector expressing an mCherry control, or a lentiviral vector expressing SMN∆7A (S270A). At 4 weeks of differentiation, there was no difference in motor neuron survival between control and SMA iPSC motor neuron cultures in any of the treatment groups. However, at 6 weeks, SMI-32 positive motor neurons showed selective loss in SMA iPSC motor neuron cultures in the untreated and lenti-mCherry groups compared to control iPSC motor neuron cultures. In contrast, lenti-SMN∆7A expression fully protects SMA iPSC-derived motor neurons. Representative images of control and SMA iPSC-derived motor neurons labeled with SMI-32 (green) and mCherry (red). Nuclei are stained with DAPI and shown in blue. *p<0.05 by ANOVA. NS = not significant. n=3

As mentioned above, experiments in an SMN-deficient chicken DT40 cell line showed that expression of SMNΔ7A, but not SMN∆7S, rescued cellular proliferation (Cho and Dreyfuss 2010). These results suggest that, when stable, SMN∆7 is intrinsically functional. To examine SMNΔ7A functionality in a more disease-relevant cell type, control and SMA induced pluripotent stem cell (iPSC) motor neuron cultures were transduced with lentiviral vectors expressing an mCherry control protein or SMNΔ7A (Fig. 6B). At 4 weeks post-differentiation, no statistical difference was observed between control and SMA motor neurons, however by 6 weeks, SMA motor neuron numbers had decreased significantly to approximately 7% of the total cell population (Fig. 6B). In contrast, expression of SMNΔ7A maintained motor neuron numbers to approximately the same level as the controls, and nearly two-fold greater than untreated cells (Fig. 6B). Thus expression of SMN∆7A improves survival of human iPSCs when differentiated into motor neuron lineages.

### SMN∆7A is a protective modifier of intermediate SMA mouse phenotypes

To examine the importance of the Slmb degron in a mammalian organismal system, two previously developed SMA mouse models were utilized. As mentioned above, the ‘Delta7’ mouse (*Smn*^*−/−*^*;SMN2;SMNΔ7*), is a model of severe SMA (Le et al. 2005), and affected pups usually die between P10 and P18 (Avg. P15). The ‘2B/−‘ mouse (*Smn*^*2B/−*^) is a module of intermediate SMA (Bowerman et al. 2012; Rindt et al. 2015) and these animals survive much longer before dying, typically between P25 and P45 (Avg. P32). Adeno-associated virus serotype 9 (AAV9) was selected to deliver the SMN cDNA isoforms to these SMA mice, as this vector has previously been shown to enter and express in SMA-relevant tissues and can dramatically rescue the SMA phenotype when expressing the wild-type SMN cDNA (Foust et al. 2010; Passini et al. 2010; Valori et al. 2010; Dominguez et al. 2011; Glascock et al. 2012).

Delivery of AAV9-SMNΔ7A at P1 significantly extended survival in the intermediate 2B/– animals, resulting in 100% of the treated pups living beyond 100 days, similar to the results obtained with the full-length AAV9-SMN construct (Fig. 7A). In contrast, untreated 2B/– animals lived, on average, only 30 days. Mice treated with AAV9-SMNΔ7S survived an average of 45 days (Fig. 7A). Mice treated with AAV9-SMNΔ7D, a phosphomimetic of the wild-type serine 270 residue, have an average life span that is equivalent or slightly shorter than that of untreated 2B/– mice (Fig. 7A). These results not only highlight the specificity of the S270A mutation in conferring efficacy to SMN∆7, but also illustrate that AAV9-mediated delivery of protein alone does not improve the phenotype.

**Figure 7:**
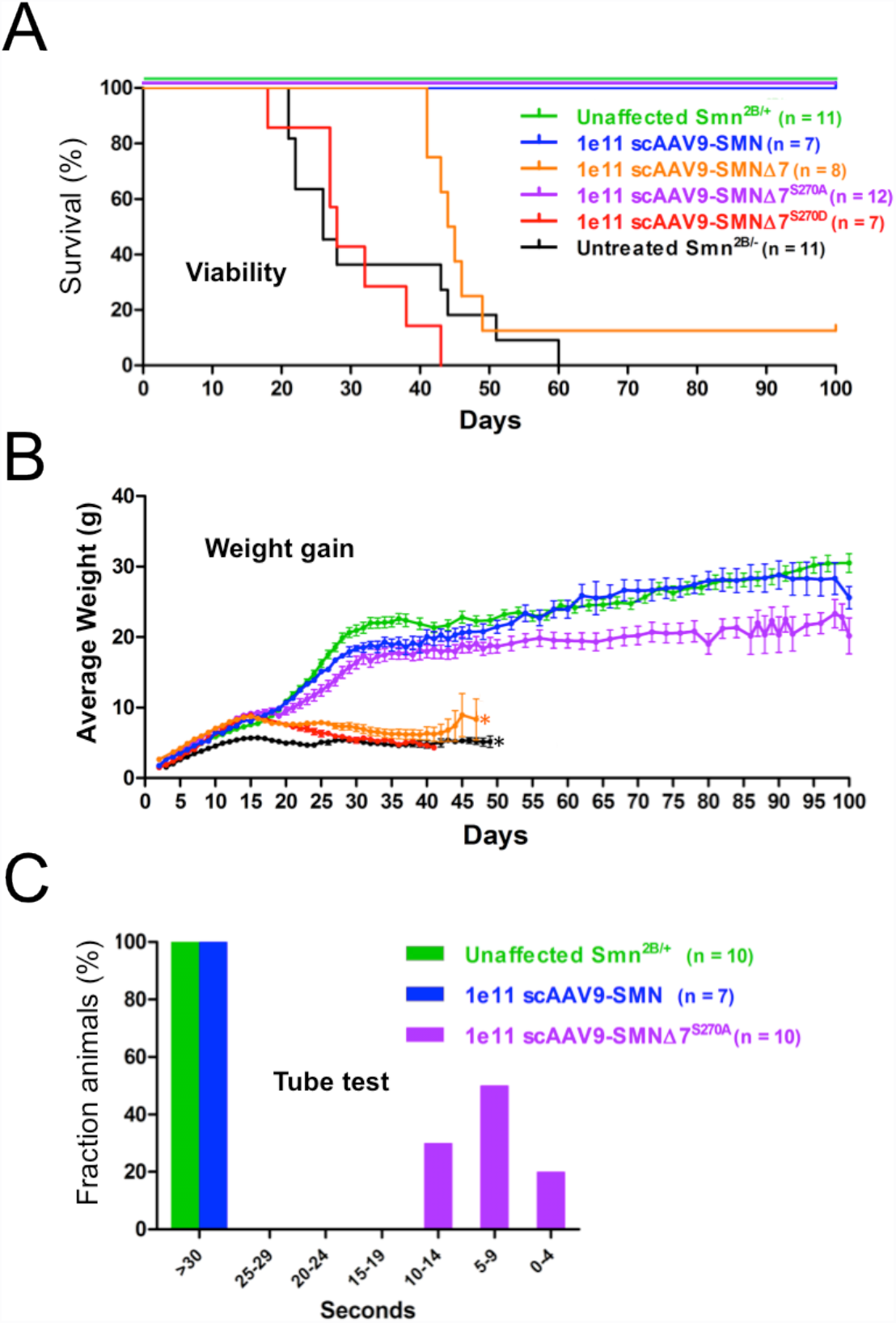
SMN∆7A is a protective modifier of intermediate SMA phenotypes in mice. *A.* Mouse genotypes include control unaffected *Smn*^*2B/+*^ mice. which have a wild-type *Smn* allele, *Smn*^*2B/+*^ (2B/—) mice treated with SCAAV9 expressing different versions of SMN, and untreated 2B/- mice, which are an intermediate mouse model of SMA. 1 e11 denotes the viral dose. scAAV9-SMN expresses full-length SMN, scAAV9-SMN∆7^S270A^ expresses truncated SMN, scAAV9-SMN∆7^S270D^ expresses truncated SMN with the S to A change in the degron, and scAAV9-SMN∆7^S270D^ expressestruncated SMN with a phosphomimic in the degron. Delivery of AAV9-SMN∆7Aat P1 significantly extended survival in the intermediate 2B/- animals, resulting in 100% of the treated pups living beyond 100 days, similar to the results obtained with the full-length AAV9-SMN construct. Untreated 2B/- animals lived, on average, only 30 days. Mice treated with AAV9-SMN∆7S survived an average of 45 days. Mice treated with AAV9 expressing SMN∆7D had an average life span equivalent or slightly worse than that of untreated 2B/- mice. *B.* Average weight (measured over time) of the animals used in panel A. AAV9-SMN∆7A treated mice also gained significantly more weight than either untreated or AAV-SMN∆7S treated animals, nearly achieving the same weight as 2B/- pups treated with full-length SMN cDNA. **C.** Mouse genotypes include control unaffected *Smn*^*2B/+*^ mice. which cany a wild-type *Smn* allele, and 2B/- mice treated with scAAV9 expressing different versions of SMN. scAAV9-SMN expresses full-length SMN and scAAV9-SMN∆7^S270A^ expressestruncated SMN with the S to A change in the degron. AAV-SMN∆7A treated animals retained their improved strength and gross motor functions at late time points (P100), as measured by their ability to splay their legs and maintain a hanging position using a modified tube-test.

We also analyzed the effects of SMN∆7A expression in the severe Delta7 mouse model (Le et al. 2005). Treatment with AAV9-SMN∆7A had only a very modest effect on Delta7 mice, as none of the animals (treated or untreated) survived weaning (Fig. S4). These findings are similar to the results in *Drosophila*. Transgenic expression of SMN∆7A in the *Smn* null background is not sufficient to rescue larval lethality (Fig. S3). Thus expression of SMN∆7A provides a clear protective benefit to the viability of intermediate mice, but not to severe SMA models.

Consistent with the lifespan data, AAV9-SMNΔ7A treated 2B/– mice gained significantly more weight than either untreated or AAV-SMNΔ7S treated animals, nearly achieving the same weight as pups treated with full-length AAV-SMN (Fig. 7B). Treatment with full-length SMN cDNA resulted in animals that were clearly stronger and more mobile, consistent with the weight data (Fig. 7C). Although they did not perform as well as mice treated with full-length SMN cDNA, the SMNΔ7A treated animals retained strength and gross motor function at late time points (e.g. P100), as measured by their ability to splay their legs and maintain a hanging position using a modified tube-test, (Fig. 7C). Animals treated with AAV9-SMN∆7D and -SMN∆7S did not survive long enough for testing.

### SCF^Slmb^ primarily targets unstable SMN monomers

As indicated in Fig. 8, our findings suggest a model whereby SMN and SMN∆7 degradation is mediated by SCF^Slmb^, a multi-component E3 ubiquitin ligase composed of Slmb, SkpA, Cul1, and Roc1 (Zheng et al. 2002, Jiang and Struhl 1998, Patton et al. 1998a; Patton et al. 1998b). Our work demonstrates that B-TrCP/Slmb binds directly to SMN (Fig. 2) and is one of a growing number of E3 ligases in the cell that can target SMN protein (Kwon et al. 2013; Han et al. 2016). SMN monomers, such as those created in SMNΔ7, are the primary targets for degradation. As shown in the model, partially-active SMN•SMN∆7 dimers and active SMN oligomers are also substrates, but to a lesser extent.

**Figure 8:**
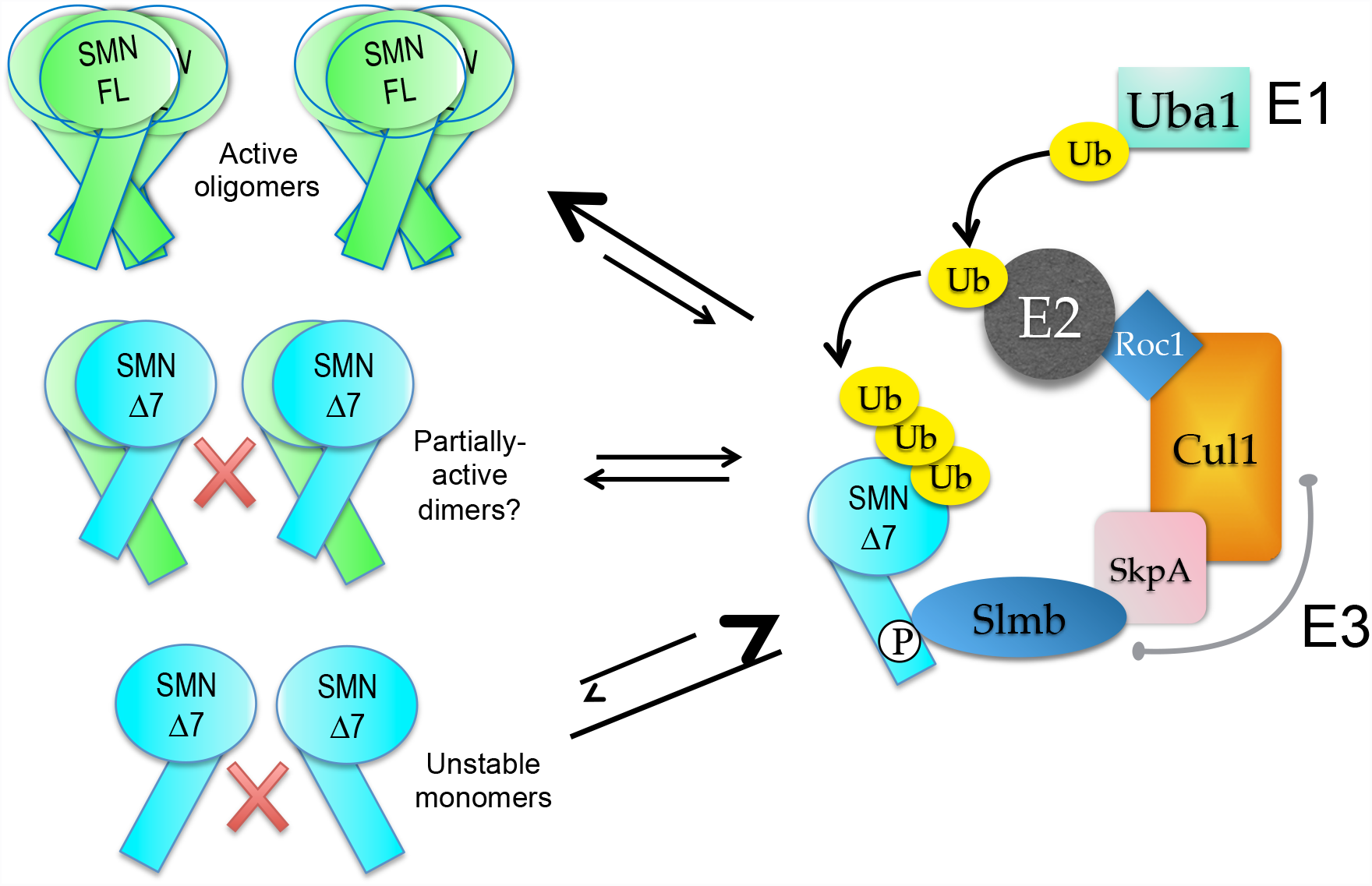
Proposed model of SMN as a substrate of SCF^slmb^ E3 ubiquitin ligase. Unstable SMN monomers, such as those created in SMN∆7, are the primary substrates for degradation. Active oligomers and partially active dimers (Praveen et al. 2014; Gupta et al. 2015) would be targeted to a lesser extent. SCF^slmb^ is a multi-component E3 ubiquitin ligase composed of Slmb, SkpA, Cull, and Rod (Zheng et al. 2002, Jiang and Struhl 1998, Patton et al. 1998a; Patton et al. 1998b). This E3 ligase works with El and E2 proteins in the ubiquitin proteasome system (UPS) to tag proteins for degradation by linkage to ubiquitin (Petroski 2008).

## Discussion

Factors that recognize the putative SMN∆7-specific degron have not been identified and the molecular mechanisms governing proteasomal access to SMN and SMN∆7 remain unclear. In this study, we isolated factors that co-purifiy with SMN from *Drosophila* embryos that exclusively express Flag-SMN. This approach reduces potential bias towards SMN partner proteins that may be more abundant in a given tissue or cell line (Charroux et al. 1999; Meister et al. 2001; Pellizzoni et al. 2002; Kroiss et al. 2008; Trinkle-Mulcahy et al. 2008; Guruharsha et al. 2011). Here, we identify the SCF^Slmb^ E3 ubiquitin ligase complex as a novel SMN binding partner whose interaction is conserved in human. Depletion of Slmb or B-TrCP by RNAi resulted in an increase in steady-state SMN levels in *Drosophila* and human cells, respectively. Furthermore, we detected four distinct high molecular weight SMN bands following immunoprecipitation with Flag-Slmb, likely corresponding to ubiquitylated isoforms. Finally, we showed that ectopic expression of SMN∆7^S270A^, but not SMN∆7 or SMN∆7^S270D^, a phosphomimetic, is a protective modifier of SMA phenotypes in animal models and human iPSC cultures.

### The SCF^Slmb^ degron is exposed by *SMN2* exon skipping

A previous study posited that a phospho-degron was specifically created by exon 7 skipping and that this event represented a key aspect of the SMA disease mechanism (Cho and Dreyfuss 2010). Our identification of a putative Slmb binding site located in the C-terminal self-oligomerization domain of *Drosophila* and human SMN has allowed us to explore the molecular details of this hypothesis. The mutation of a conserved serine within the Slmb degron not only disrupted the interaction between SMN and Slmb, but also stabilized full-length SMN and SMN∆7. Notably, the degron mutation has a greater effect on SMN levels (both full-length and

∆7) when made in the context of a protein that does not efficiently self-oligomerize. These and other findings strongly suggest that the Slmb degron is uncovered when SMN is monomeric, whereas it is less accessible when SMN forms higher-order multimers. On the basis of these results, we conclude that *SMN2* exon skipping does not <u>*create*</u> a potent protein degradation signal; rather, it <u>*exposes*</u> an existing one.

### SMN targeting by multiple E2 and E3 systems

SMN degradation via the UPS is well-established (Chang et al. 2004; Burnett et al. 2009; Kwon et al. 2011). Using candidate approaches, investigators have studied other E3 ligases that have been reported to target SMN for degradation in cultured human cells (Han et al. 2016; Hsu et al. 2010; Kwon et al. 2013). Given our findings, it is therefore likely that SMN is targeted by multiple E3 ubiquitin ligases, as this regulatory paradigm has been demonstrated for a number of proteins (e.g. p53; Jain and Barton 2010). Targeting of a single protein by multiple E3 ligases is thought to provide regulatory specificity by expressing the appropriate degradation complexes only within certain tissues, subcellular compartments or developmental timeframes. Moreover, ubiquitylation does not always result in immediate destruction of the target; differential use of ubiquitin lysine linkages or chain length can alter a protein’s fate (Mukhopadhyay and Riezman 2007; Ikeda and Dikic 2008; Liu and Walters 2010).

Avenues of future exploration include determination of the E2 proteins that partner with SCF^Slmb^ as well as the types of ubiquitin lysine chain linkages they add to SMN. These two questions are interconnected, as ubiquitin linkage specificity is determined by the E2 (Ye and Rape 2009). Lysine 48 (K48) linked chains typically result in degradation of the targeted protein by the 26S proteasome, whereas lysine 63 (K63) linkage is more commonly associated with lysosomal degradation and nonproteolytic functions such as endocytosis (Tan et al. 2007; Kirkin et al. 2009; Lim and Lim 2010). Interestingly, recent work has implicated defects in endocytosis in SMA (Custer and Androphy 2014; Dimitriadi et al. 2016; Hosseinibarkooie et al. 2016). It remains to be determined how the ubiquitylation status of SMN might intersect with endocytic functions.

### Does SMN function as a signaling hub?

In the Flag-SMN pulldown, we identified three E2 proteins as potential SMN interacting partners (Fig. 1C). Among them, Bendless (Ben) is particularly interesting. Ben physically interacts with TRAF6, an E3 ligase that functions together with Ube2N/Ubc13/Ben in human cells (Kim and Choi 2017). TRAF6 is an activator of NF-kB signaling, and its interaction with SMN is thought to inhibit this activity (Kim and Choi 2017). Notably, Ube2N/Ben heterodimerizes with Uev1a to form K63 ubiquitin linkages on target proteins (Ye and Rape 2009; van Wijk and Timmers 2010; Komander and Rape 2012; Marblestone et al. 2013; Zhang et al. 2013). Furthermore, Ben-Uev1a is involved in upstream activation of both JNK and IMD signaling in *Drosophila* (Paquette et al. 2010; Zhou et al. 2005). Previously, we and others have shown that JNK signaling is dysregulated in animal models of SMA (Garcia et al. 2013; Genabai et al. 2015; Garcia et al. 2016; Ahmad et al. 2016). Moreover, mutations in all three components of SCF^Slmb^ lead to constitutive expression of antimicrobial peptides, which are also downstream of the IMD pathway (Khush et al. 2002). Together, these findings suggest the interesting possibility of SMN functioning as a signaling hub that links the UPS to the JNK and IMD pathways, all of which have been shown to be disrupted in SMA.

### Phosphorylation of the Slmb degron within SMN

As Slmb is known to recognize phospho-degrons, one of the first questions raised by our study concerns the identity of the kinase(s) responsible for phosphorylating the degron in SMN. A prime candidate is GSK3β, as this kinase recognizes a motif (SxxxS/T; Liu et al. 2007; Lee et al. 2013) that includes the degron and extends N-terminally (^262^SxxxSxxxSxxxT^274^, numbering as per human SMN). In support of this hypothesis, we identified the *Drosophila* GSK3β orthologue, Shaggy (Sgg), in our SMN pulldowns (Fig. 1C). Moreover, GSK3β inhibitors as well as siRNA mediated knockdown of GSK3β were shown to increase SMN levels, primarily by stabilizing the protein (Makhortova et al. 2011; Chen et al. 2012). Finally, GSK3β is also responsible for phosphorylation of a degron in β-catenin, a well-characterized SCF^Slmb^ substrate (Liu et al. 2002). SMA mice have low levels of UBA1 (E1) ultimately leading to accumulation of β-catenin (Wishart et al. 2014). Pharmacological inhibition of β-catenin improved neuromuscular pathology in *Drosophila*, zebrafish, and mouse SMA models. β-catenin had previously been shown to regulate motor neuron differentiation and stability by affecting synaptic structure and function (Murase et al. 2002; Li et al. 2008; Ojeda et al. 2011). β-catenin also regulates motor neuron differentiation by retrograde signaling from skeletal muscle (Li et al. 2008). The connections of UBA1 and multiple SCF^Slmb^ substrates to motor neuron health thus places the UPS at the center of SMA research interest.

### Concluding remarks

In summary, this study identifies conserved factors that regulate SMN stability. To our knowledge, this work represents the first time that SMN complexes have been purified in the context of an intact developing organism. Using this approach, we have demonstrated that the SCF^Slmb^ E3 ligase complex interacts with a degron embedded within the self-oligomerization domain of SMN. Our findings establish plausible connections to disease-relevant cellular processes and signaling pathways. Further, they elucidate a model whereby accessibility of the SMN phospho-degron is regulated by self-multimerization, providing an elegant mechanism for balancing functional activity with degradation.

## Experimental procedures

### Fly stocks and transgenes

Oregon-R was used as the wild-type control. The Smn^X7^ microdeletion allele (Chang et al. 2008) was a gift from S. Artavanis-Tsakonis (Harvard University, Cambridge, USA). This deficiency removes the promoter and the entire SMN coding region, leaving only the final 44bp of the 3’ UTR. All stocks were cultured on molasses and agar at room temperature (24 ± 1°C) in half-pint bottles. The Smn transgenic constructs were injected into embryos by BestGene Inc. (Chino Hills, CA) as described in Praveen et al. 2014. In short, a ~3kb fragment containing the entire *Smn* coding region was cloned from the *Drosophila* genome into the pAttB vector (Bischof et al. 2007). A 3X FLAG tag was inserted immediately downstream of the start codon of dSMN. Point mutations were introduced into this construct using Q5 (NEB) site-directed mutagenesis according to manufacturer’s instructions. The basal Smn construct used, vSmn, contained three single amino acid changes and the addition of the MGLR motif to make fruitfly Smn more similar to the evolutionarily conserved vertebrate Smn. Subsequently generated constructs used vSmn as a template and consist of the amino acid changes detailed in Figure 4. Y203C, G206S, and G210V were previously published in Praveen et al. 2014.

### Drosophila embryo protein lysate and mass spectrometry

0-12h Drosophila embryos were collected from Oregon-R control and Flag-SMN flies, dechorionated, flash frozen, and stored at −80C. Embryos (approx. 1gr) were then homogenized on ice with a Potter tissue grinder in 5 mL of lysis buffer containing 100mM potassium acetate, 30mM HEPES-KOH at pH 7.4, 2mM magnesium acetate, 5mM dithiothreitol (DTT) and protease inhibitor cocktail. Lysates were centrifuged twice at 20000 rpm for 20min at 4C and dialyzed for 5h at 4C in Buffer D (HEPES 20mM pH 7.9, 100mM KCl, 2.5 mM MgCl_2_, 20% glycerol, 0.5 mM DTT, PMSF 0.2 mM). Lysates were clarified again by centrifugation at 20000 rpm for 20 min at 4C. Lysates were flash frozen using liquid nitrogen and stored at −80C before use. Lysates were then thawed on ice, centrifuged at 20000 rpm for 20 min at 4C and incubated with rotation with 100 µL of EZview Red Anti-FLAG M2 affinity gel (Sigma) for 2h at 4C. Beads were washed a total of six times using buffer with KCl concentrations ranging from 100mM to 250mM with rotation for 1 min at 4C in between each wash. Finally, Flag proteins were eluted 3 consecutive times with one bed volume of elution buffer (Tris 20mM pH 8, 100 mM KCl, 10% glycerol, 0.5 mM DTT, PMSF 0.2 mM) containing 250ug/mL 3XFLAG peptide (sigma). The entire eluate was used for mass spectrometry analysis on an Orbitrap Velos instrument, fitted with a Thermo Easy-spray 50cm column.

### Tissue culture and transfections

S2 cell lines were obtained from the Drosophila Genome Resource Center (Bloomington, IL). S2 cells were maintained in SF900 SFM (Gibco) supplemented with 1% penicillin/streptomycin and filter sterilized. Cells were removed from the flask using a cell scraper and passaged to maintain a density of approximately 10^6^-10^7^ cells/mL. S2 cells were transferred to filter sterilized SF900 SFM (Gibco) without antibiotics prior to transfection with Cellfectin II (Invitrogen). Transfections were performed according to Cellfectin II protocol in a final volume of 4 mL in a T-25 flask containing 10^7^ cells that were plated one hour before transfection. The total amount of DNA used in transfections was 2.5ug. Human embryonic kidney HEK-293T and HeLa cells were maintained at 37C with 5% CO_2_ in DMEM (Gibco) supplemented with 10% FBS and 1% penicillin/streptomycin (Gibco). 1×10^6^ - 2×10^6^ cells were plated in T-25 flasks and transiently transfected with 1-2ug of plasmid DNA per flask using Lipofectamine (Invitrogen) or FuGENE HD transfection reagent (Roche Applied Science, Indianapolis, IN) according to the manufacturer’s protocol. Cells were harvested 24-72 h posttransfection.

For siRNA transections, HeLa cells were plated subconfluently in T-25 flasks and transfected with 10nm of siRNA (Gift from Mike Emanuele lab) and 17uL Lipofectamine RNAi MAX (Invitrogen) in 5mL total media according to manufacturers instructions. After 48h of transfection cells were harvested. For RNAi in S2 cells using dsRNA, 10^7^ cells were plated in each well of a 6-well plate in 1 mL of media. Cells were treated ~ every 24h with 10ug/mL dsRNA targeted against Slmb, Oskar, or Gaussia Luciferaese (as controls) as described in Rogers and Rogers 2008.

### In vitro binding assay

GST and GST-SMN were purified from *E. coli*. In brief, cells transformed with BL21*GST-SMN were grown at 37˚C overnight and then induced using 1 mM IPTG. Recombinant protein was extracted and purified using Glutathione sepharose 4B beads. GST-B-TrCP1 was purchased from Novus Biologicals (cat# H00008945). SMN•Gem2 complexes were co-expressed in *E. coli* using SMN∆5 and Gemin2(12-280) constructs, as described in Gupta et al. (2015). Glutathione sepharose 4B beads were washed 3x with PBS. GST alone, GST-SMN, or GST-B-TrCP1 were attached to beads during 4h-overnight incubation at 4˚C in PBS with rotation. Beads were then washed 3x with modified RIPA buffer (50mM Tris-HCl, pH 7.5, 250 mM NaCl, 1mM EDTA, 1% NP-40). 20uL of beads with ~2ug attached GST-tagged protein (as determined by Coomassie stain with BSA standard) were added to 200uL modified RIPA buffer with 100ug/mL BSA block. 2ug of SMN•Gem2 was added and the mixture was rotated end over end at 4˚C overnight. Beads were then washed 3x with modified RIPA buffer and 10uL SDS loading buffer was added followed by boiling for 5 minutes.

### *In vivo* ubiquitylation assay

The *in vivo* ubiquitylation assay was performed as described previously (Choudhury et al. 2016). Briefly, HEK-293T cells were transfected as indicated in 10 cm dishes using Lipofectamine2000 (Thermo Fisher Scientific). The day after, cells were treated with 20 µM of MG132 for 4 hours and then harvested in PBS. 80% of the cell suspension was lysed in 6M Guanidine-HCl containing buffer and used to pull down His-Ubiquitinated proteins on Ni2+-NTA beads, while the remaining 20% was used to prepare inputs. Ni2+ pull down eluates and inputs were separated through SDS-PAGE and analyzed by western blot.

### Cycloheximide treatment

Following RNAi treatment, S2 cells were pooled, centrifuged and resuspended in fresh media. 1/3 of these cells were frozen and taken as the 0h timepoint. The remainders of the cells were replated in 6 well plates. 100ug/mL cycloheximide (CHX) was added to each sample, and cells were harvested at 2 and 6 hours following treatment.

### Immunoprecipitation

Clarified cell lysates were precleared with immune-globulin G (IgG) agarose beads for 1h at 4C and again precleared overnight at 4C. The precleared lysates were then incubated with Anti-FLAG antibody crosslinked to agarose beads (EZview Red Anti-FLAG M2 affinity gel, Sigma) for 2h at 4C with rotation. The beads were washed with lysis buffer or modified lysis buffer six times and boiled in SDS gel-loading buffer. Eluted proteins were run on an SDS-PAGE for western blotting.

### Antibodies and Western blotting

Larval and adult lysates were prepared by crushing the animals in lysis buffer (50mM Tris-HCl, pH 7.5, 150 mM NaCl, 1mM EDTA, 1% NP-40) with 1X (adults) or 10x (larvae) protease inhibitor cocktail (Invitrogen) and clearing the lysate by centrifugation at 13,000 RPM for 10 min at 4ºC. S2 cell lysates were prepared by suspending cells in lysis buffer (50mM Tris-HCl, pH 7.5, 150 mM NaCl, 1mM EDTA, 1% NP-40) with 10% glycerol and 1x protease inhibitor cocktail (Invitrogen) and disrupting cell membranes by pulling the suspension through a 25 gauge needle (Becton Dickinson). The lysate was then cleared by centrifugation at 13,000 RPM for 10 min at 4ºC. Human cells (293Ts and HeLas) were first gently washed in ice-cold 1X PBS, then collected in ice-cold 1X PBS by scraping. Cells were pelleted by spinning at 1000 rpm for 5 min. The supernatant was removed and cells were resuspended in ice cold lysis buffer (50mM Tris-HCl, pH 7.5, 150 mM NaCl, 1mM EDTA, 1% NP-40) and allowed to lyse on ice for 30 min. After lysing, the lysate was cleared by centrifuging the cells for 10 min at 13000 at 4C. Western blotting on lysates was performed using standard protocols. Rabbit anti-dSMN serum was generated by injecting rabbits with purified full-length dSMN protein (Pacific Immunology Corp, CA), and was subsequently affinity purified. For Western blotting, dilutions of 1 in 2,500 for the affinity purified anti-dSMN, 1 in 20,000 (fly) or 1 in 5,000 (human) for anti-α tubulin (Sigma), 1 in 10,000 for monoclonal anti-Flag (Sigma), 1 in 1,000 for anti-Slmb (gift from Greg Rogers), 1 in 2,500 for anti-human SMN (BD Biosciences), 1 in 1,000 for anti-B-TrCP (gift from MB Major lab), 1 in 10,000 for polyclonal anti-Myc (Santa Cruz), and 1 in 2,000 for anti-GST (abcam) were used.

### Larval locomotion

*Smn* control and mutant larvae (73-77 hours post egg-laying) were placed on a 1.5% agarose molasses tray five at a time. The tray was then placed in a box with a camera and the larvae were recorded moving freely for 60 seconds. Each set of larvae was recorded three times, and one video was chosen for analysis based on video quality. The videos were then converted to AVI files and analyzed using the wrMTrck plug-in of the Fiji software. The "Body Lengths per Second" was calculated in wrMTrck by dividing the track length by half the perimeter and time (seconds). P-values were generated using a multiple comparison ANOVA.

### SMA Mouse Models

Two previously developed SMA mouse models were utilized. The ‘Delta7’ mouse (*Smn*^−/−^;*SMN2;SMNΔ7*), is a model of severe SMA (Le et al. 2005). The ‘2B/–‘ mouse (*Smn*^*2B/–*^) is a model of intermediate SMA (Bowerman et al. 2012; Rindt et al. 2015). Adeno-associated virus serotype 9 (AAV9) delivered SMN cDNA isoforms to these SMA mice, as previously described (Foust et al. 2010; Passini et al. 2010; Valori et al. 2010; Dominguez et al. 2011; Glascock et al. 2012). Gross motor function was measured using a modified tube-test which tests the ability of mice to splay their legs and maintain a hanging position.

### Human iPSC Cell culture

Human iPSCs from two independent unaffected control and two SMA patient lines were grown as pluripotent colonies on Matrigel substrate (Corning) in Nutristem medium (Stemgent). Colonies were then lifted using 1mg/ml Dispase (Gibco) and maintained as floating spheres of neural progenitor cells in the neural progenitor growth medium Stemline (Sigma) supplemented with 100ng/ml human basic fibroblast growth factor (FGF-2, Miltenyi), 100ng/ml epidermal growth factor (EGF, Miltenyi), and 5µg/ml heparin (Sigma-Aldrich) in ultra-low attachment flasks. Aggregates were passaged using a manual chopping technique as previously described (Svendsen et al. 1998; Ebert et al. 2013). To induce motor neuron differentiation, neural progenitor cells were cultured in neural induction medium (1:1 DMEM/F12 (Gibco), 1x N2 Supplement (Gibco), 5µg/mL Heparin (Sigma), 1x Non-Essential Amino Acids (Gibco), and 1x Antibiotic-Antimycotic (Gibco)) plus 0.1µM all-trans retinoic acid (RA) for two weeks; 1µM Purmorphamine (PMN, Stemgent) was added during the second week. Spheres were then dissociated with TrypLE Express (Gibco) and plated onto Matrigel-coated 12mm coverslips in NIM plus 1µM RA, 1µM PMN, 1x B27 Supplement (Gibco), 200ng/mL Ascorbic Acid (Sigma), 1µM cAMP (Sigma), 10ng/mL BDNF (Peprotech), 10ng/mL GDNF (Peprotech)). One week post-plating, cells were infected with lentiviral vectors (MOI = 5) expressing mCherry alone or SMN S270A-IRES-mCherry. Transgenes in both viruses were under the control of the EF1α promoter. Uninfected cells served as controls. Cells were analyzed at 1 and 3 weeks post-infections, which was 4 and 6 weeks of total differentiation (Ebert et al. 2009; Sareen et al. 2013).

### Immunocytochemistry

Coverslips were fixed in 4% paraformaldehyde (Electron Microscopy Sciences) for 20 minutes at room temperature and rinsed with PBS. Cells were blocked with 5% Normal Donkey Serum (Millipore) and permeabilized in 0.2% TritonX-100 (Sigma) for 30 minutes at room temperature. Cells were then incubated in primary antibody solution for 1 hour, rinsed with PBS, and incubated in secondary antibody solution for 1 hour at room temperature. Finally, nuclei were labeled with Hoechst nuclear stain (Sigma) to label DNA and mounted onto glass slides using FluoroMount medium (SouthernBiotech). Primary antibodies used were mouse anti-SMI-32 (Covance SMI-32R, 1:1000) and rabbit anti-mCherry (ThermoFisher, 1:1000). Secondary antibodies used were donkey anti-rabbit Cy3 (Jackson Immunoresearch 711-165-152) and donkey anti-mouse AF488 (Invitrogen A21202).

### Immunocytochemical Analysis

Images were acquired from five random fields per coverslip using an inverted fluorescent microscope (Nikon) and NIS Elements software. Images were blinded and manually analyzed for antigen specificity with NIS Elements software.

## Acknowledgements

This work was supported by NIGMS R01 GM118636 (to A.G.M.). K.M.G. was supported by graduate research fellowship DGE-1144081 from the NSF. Work in the Wagner lab (D.B and E.J.W.) was supported by the Welch Foundation (H1889). We also thank the Peifer, Major, and Rogers laboratories for reagents, advice and expertise.

---

**Figure S1:**
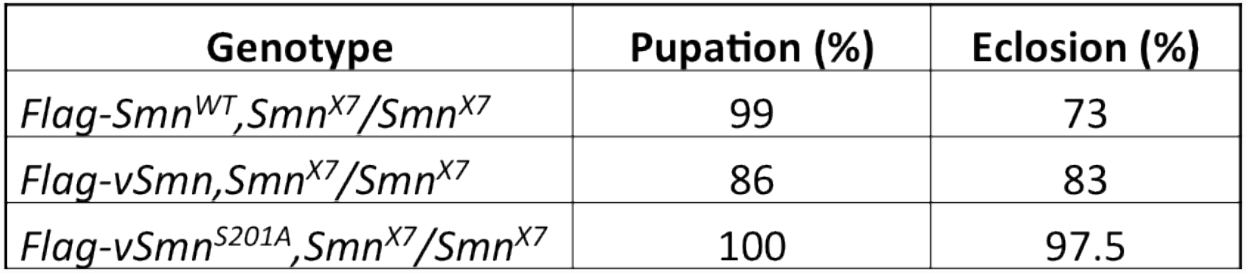
A) Transgenic flies expressing Flag-vSmn and Flag-vSmn^S201A^ in the background of an *Smn*^*X7*^ null mutation are fully viable. The eclosion frequencies of these animals are consistently higherthan those that express Flag-Smn^WT^ in the background of an *Smn*^*X7*^ null mutation. The data for each genotype are expressed as a fraction of pupae or adults over the total number of starting larvae, n=200.

**Figure S2:**
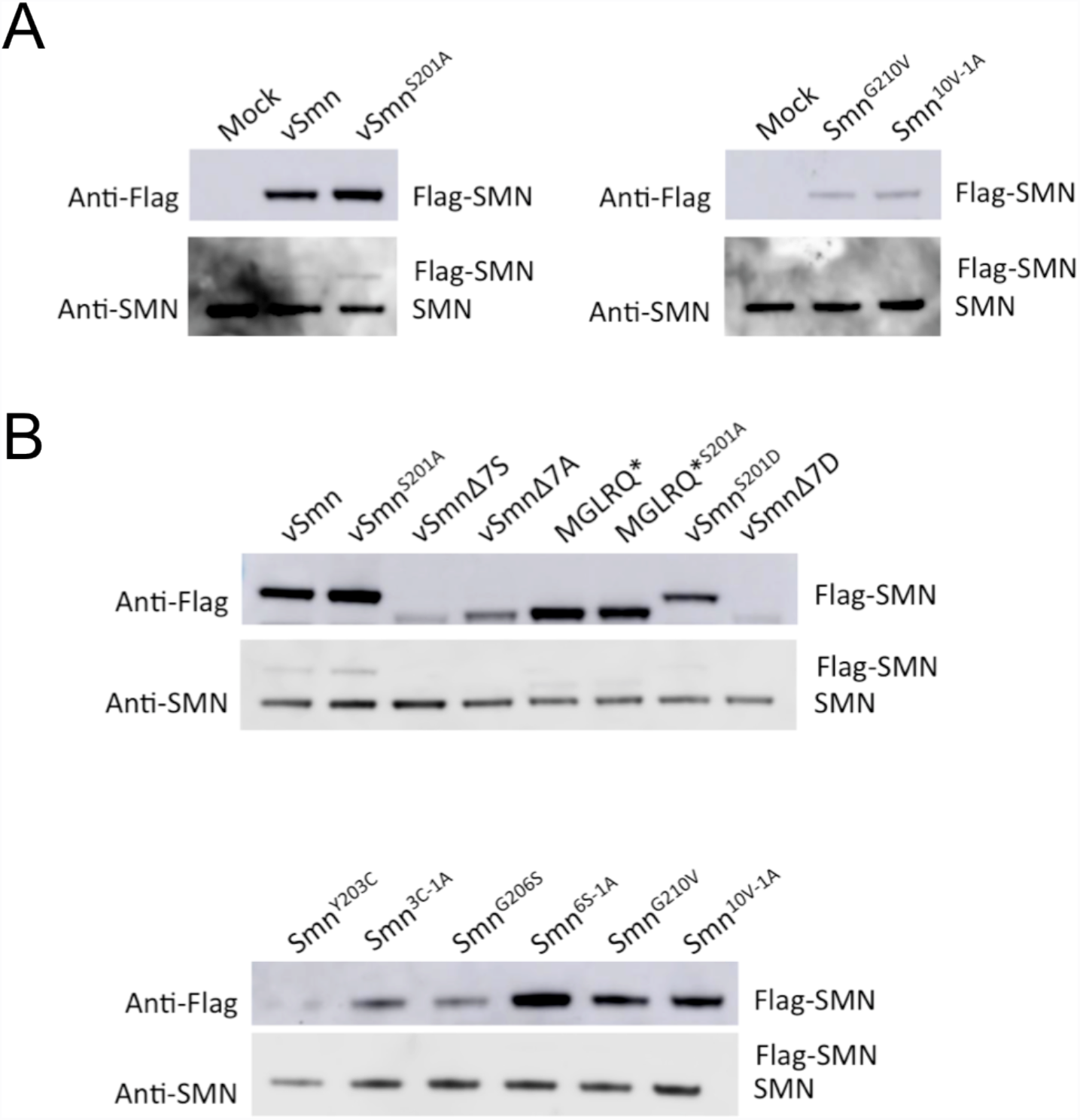
A) The expression of endogenous SMN in S2 cells following transient transfection of either modified ‘vertebrate’ SMN constructs (vSMN and vSMN^S201A^) or Drosophila SMN constructs (Smn^G210V^ and Smn^10V-1A^) is unaffected, as compared to mock transfection. Protein is detected by anti-Flag antibody or anti-SMN antibody as indicated to the left of the blots. B) Transient transfections in S2 cells express Flag-SMN from the endogenous promoter. Protein levels of all transfected constructs are lower than endogenous SMN protein levels. Protein is detected by anti-Flag antibody or anti-SMN antibody as indicated to the left of the blots.

**Figure S3:**
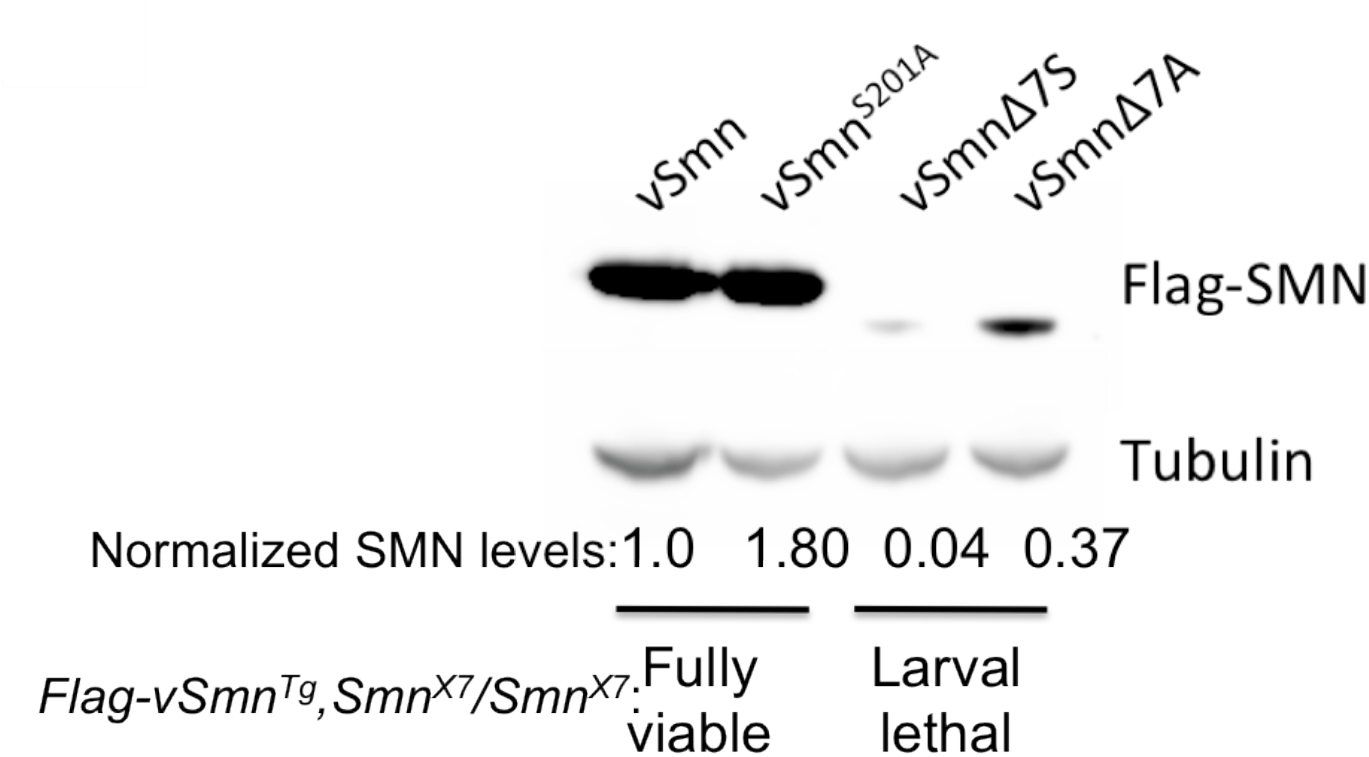
Western blotting was used to determine protein levels of each of the SMN constructs, with expression driven by the endogenous promoter, in transgenic adult flies. Protein lysates were made by pooling 40-50 adult flies. Flies with the following genotypes were analyzed in this experiment: *Flag-vSmn,Smn*^*X7*^*/Tm6b* (vSmn), *Flag-vSmn*^S201A^,*Smn*^X7^*/Tm6b* (vSmn^s201A^), *Flag-vSmn∆7S,Smn*^*X7*^*/Tm6b* (vSMN∆7S) or *Flag-vSMN∆7A,Smn*^*X7*^*/Tm6b* (vSMN∆7A). Both the vSMN and VSMN∆7S proteins show increased levels when the serine is mutated to an alanine, indicating disruption of the normal degradation of SMN. Flag-SMN was detected using anti-Flag antibody. Normalized fold change as compared to vSmn levels is indicated at the bottom.

**Figure S4:**
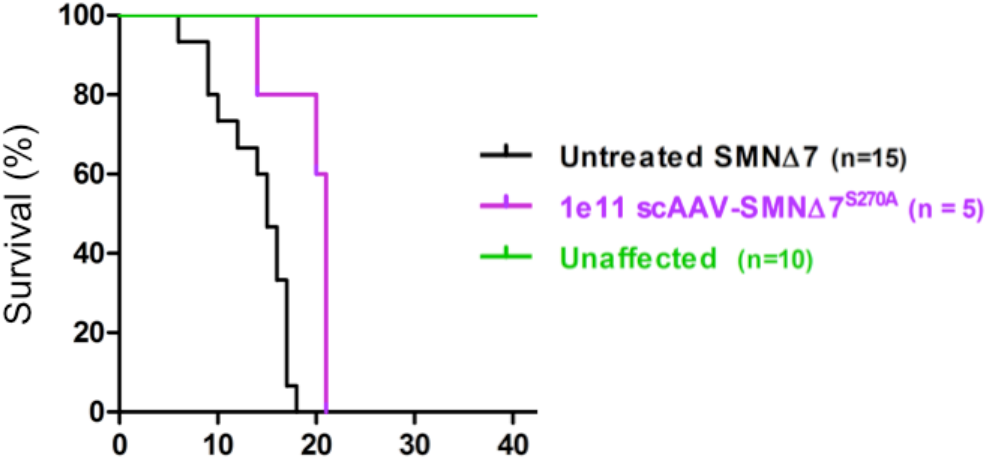
Survival analysis of the effects of SMN∆7A expression in the severe Delta7 mouse model. Genotypes include untreated SMN∆7 mice, which are a severe mouse model of SMA, SMN∆7 mice treated with scAAV9 expressing SMN∆7^S270A^, truncated SMN with the S to Achange in the degron, and control unaffected mice, which have a wild-type Smn allele. Treatment with AAV9-SMN∆7A had only a very modest effect on viability and none of the animals survived weaning. 1e11 denotes the viral dose.

